# Evolutionary mechanisms that determine which bacterial genes are carried on plasmids

**DOI:** 10.1101/2020.08.04.236455

**Authors:** Sonja Lehtinen, Jana S. Huisman, Sebastian Bonhoeffer

**Affiliations:** Institute for Integrative Biology, Department of Environmental System Science, ETH Zürich, Zürich, Switzerland; Swiss Institute of Bioinformatics, Lausanne, Switzerland

## Abstract

The evolutionary pressures that determine the location (chromosomal or plasmid-borne) of bacterial genes are not fully understood. We investigate these pressures through mathematical modelling in the context of antibiotic resistance, which is often found on plasmids. Our central finding is that gene location is under positive frequency-dependent selection: the higher the frequency of one form of resistance compared to the other, the higher its relative fitness. This can keep moderately beneficial genes on plasmids, despite occasional plasmid loss. For these genes, positive frequency-dependence leads to a priority effect: whichever form is acquired first – through either mutation or horizontal gene transfer – has time to increase in frequency and thus become difficult to displace. Higher rates of horizontal transfer of plasmid-borne than chromosomal genes therefore predict moderately beneficial genes will be found on plasmids. Gene flow between plasmid and chromosome allows chromosomal forms to arise, but positive frequencydependent selection prevents these from establishing. Further modelling shows this effect is particularly pronounced when genes are shared across a large number of species, suggesting antibiotic resistance genes are often found on plasmids because they are moderately beneficial across many species. We also revisit previous theoretical work – relating to the role of local adaptation in explaining gene location and to plasmid persistence – in light of our findings.

**Impact Statement:** Bacterial genes can either reside on the chromosome or on plasmids, extra-chromosomal genetic structures which can be transferred from cell to cell. The distribution of genes between plasmid and chromosome is not random: certain types of genes are particularly likely to be plasmid-associated. This includes a number of clinically important traits, such as antibiotic resistance and virulence factors. The evolutionary mechanisms that give rise to this pattern are not well understood. Plasmids are occasionally lost during cell replication and thus less reliably inherited than the chromosome, and genes are free to transition between plasmid and chromosome: so what keeps genes on plasmids? We address this question using a mathematical model. The key prediction from our model is that the relative fitness of chromosomal and plasmid-borne genes depends on their relative frequencies (‘positive frequency-dependent selection’). In other words, the fitness of a plasmid-borne gene will be higher in a population in which the chromosomal gene is rare (and vice-versa). This positive-frequency dependence can keep moderately beneficial genes on plasmids, despite occasional plasmid loss. This leads to a priority effect: whichever form of the gene (i.e. plasmid-borne or chromosomal) is acquired first has time to increase in frequency and thus become difficult to displace. Therefore, the relative rate of acquiring the gene on the plasmid vs the chromosome predicts where the gene will be found. Further modelling shows this effect is particularly pronounced when genes are beneficial across a large number of species. All together, the hypothesis that emerges from our work is that plasmid-borne genes are moderately beneficial; functional across a large number of species; and rarely acquired through chromosomal mutation. We suggest traits like antibiotic resistance are often found on plasmids because these genes commonly fulfill these criteria.

## Introduction

Plasmids are extra-chromosomal genetic structures that can replicate independently from the chromosome and be transferred from cell-to-cell. These structures play a key role in bacterial evolution: in addition to genes that enable their own replication and spread, plasmids can carry genes that are beneficial to the bacterial host. This makes plasmids an important vehicle of horizontal gene transfer, both within and between species.

There is evidence that the types of genes found on plasmids differ from those found on the chromosome. In general terms, plasmid-borne genes are part of the accessory rather than core genome [1, 2]. Furthermore, some accessory functions appear to be particularly likely to be plasmid associated [2, 3, 4] -these include virulence factors [5], antibiotic resistance [6, 7], heavy metal tolerance [8], and bacteriocins (toxins involved in inter-strain competition) [9].

These patterns are intriguing: genes can transition between plasmids and the chromosome [10]; thus, over evolutionary time-scales, genes are likely to have experienced both plasmid and chromosomal backgrounds [3]. There must therefore be evolutionary mechanisms that give rise to the association between plasmids and particular gene functions. Understanding these mechanisms is important, both because of the role plasmids play in bacterial evolution and because of the clinical relevance of plasmid-associated functions.

The absence of core genes from plasmids is relatively well understood. Firstly, core genes are more likely to be highly connected in cellular networks [11], thus making them less transferable across genetic backgrounds (‘complexity hypothesis’ [12]). Secondly, inheritance of plasmids is less stable than inheritance of chromosomes: daughter cells do not always inherit a copy of the plasmid during cell division (‘segregation loss’). Thus, if a gene is essential to the survival of the cell, we would expect it to locate onto the chromosome. Plasmids encode mechanisms to prevent segregation loss [13] and plasmid loss is therefore rare [14], but even low rates of plasmid loss are predicted to make essential genes chromosomal rather than plasmid-borne [15]. Thirdly, horizontal gene transfer can allow lineages to recover lost genes [16, 17, 18], but this mechanism is less relevant to essential genes, as lineages without these genes would themselves be lost [19].

The over-representation of particular functions on plasmids remains puzzling. In the specific case of cooperative traits, one possible explanation is that mobility is beneficial because it enforces the cooperation of neighbouring cells [20, 21, 2]. More generally, it has been suggested that plasmid-borne genes code for local adaptation: if a gene is beneficial in a specific environment, being plasmid-borne would allow this gene to spread into immigrant lineages lacking the trait, thus maintaining the gene on the plasmid [3, 16]. A related suggestion is that temporal fluctuation in selection pressure would favour plasmid-borne genes: being plasmid-borne would allow the gene to increase in frequency faster during periods of positive selection and to persist through plasmid transfer during period of negative selection [6, 22].

The local adaptation explanation is based on the assumption that the immigrant lineage is susceptible to infection by the local plasmid, i.e. that it does not already carry the plasmid (either with or without the locally beneficial gene). This assumption arises from estimates of plasmid transfer rates in liquid culture, which suggest plasmids are not transmissible enough, relative to their cost and the rate of plasmid loss, to exists as pure parasites (i.e. without carrying genes beneficial for the host). Thus, outside the local niche with the beneficial gene, plasmids would not persist [16].

More recently however, this assumption has been called into question [23]. Estimates of plasmid transfer rates are higher in biofilms than in liquid culture [24] and there is evidence that compensatory evolution acts to ameliorate plasmid cost [25, 26], particularly in the presence of spatially heterogeneous selection [27]. Indeed, recent experimental evidence suggests conjugative plasmids can persist through plasmid transfer despite not being beneficial [28]. Furthermore, genomic studies detect the same plasmid backbones with different gene content (e.g. with and without particular resistance genes [10]). Taken together, this evidence suggests models of plasmid dynamics should include the possibility of both gene-bearing and gene free versions of the plasmid.

We therefore revisit the question of why some genes are carried on plasmids, accounting for competition between plasmids, as well as chromosomes, with and without the focal gene. This yields a relatively complex model, which is difficult to study analytically (though previous work has addressed such a model through simulation [6]). Here, we adopt an approach which makes the otherwise analytically unsolvable model mathematically tractable. For readability, our model is framed in terms of antibiotic resistance genes, though our results are more broadly applicable. Through a combination of analysis and simulation, we find that, indeed, in presence of the non-beneficial plasmid, local adaptation does not explain why some genes are plasmid-borne. However, gene location is under positive frequency-dependent selection, which leads to a priority effect: whichever form is acquired first has time to increase in frequency and thus become difficult to displace. The presence of resistance genes on plasmids could therefore be explained by a greater rate of acquisition of plasmid-borne than chromosomal resistance.

## Results

### Model

We consider a population consisting of cells which can be either chromosomally resistant or chromosomally sensitive (*R* or *S*), with no plasmid, a resistant plasmid, or a sensitive plasmid (∅, *R*, or *S*). This gives rise to six possible cell types: *R*∅, *RR, RS, S*∅, *SR* and *SS* (first letter denotes the chromosome, the second letter the plasmid).

We begin by developing a model of the dynamics of these cells (Figure 1) and denote the density of each cell type by *N*_•_. Cells replicate at rate *λ*. Competition between cells is captured through a density-dependent death rate *γT*, where *T* is the total cell density (*T* = *N*_*S*∅_ + *N*_*SS*_ + *N*_*SR*_ + *N*_*R*∅_ + *N*_*RS*_ + *N*_*RR*_). Plasmids spread through density-dependent transmission between cells at rate *β* and are lost during cell replication with probability *s* (‘segregation loss’). Plasmid carriage is associated with a fitness cost *c*_*P*_, which reduces the replication rate *λ* by a factor of 1 − *c*_*P*_. We assume cells can only be infected with one plasmid at a time. Cells with no resistance (*S*∅ and *SS*) experience an additional death rate *A* from antibiotic exposure. Resistance is associated with a fitness cost, which reduces the replication rate by a factor of 1 − *c*_*R*_. We assume both the fitness cost and the effectiveness of resistance are the same whether resistance is chromosomal or plasmid-borne. Cells which have both chromosomal and plasmid-borne resistance experience a dual fitness cost (1 − *c*_*R*_)^2^. The effect of modifying these assumptions is explored in the Supporting Information (SI Section 2). Our main results are generally robust, with sensitivities highlighted in the main text.

**Figure 1:**
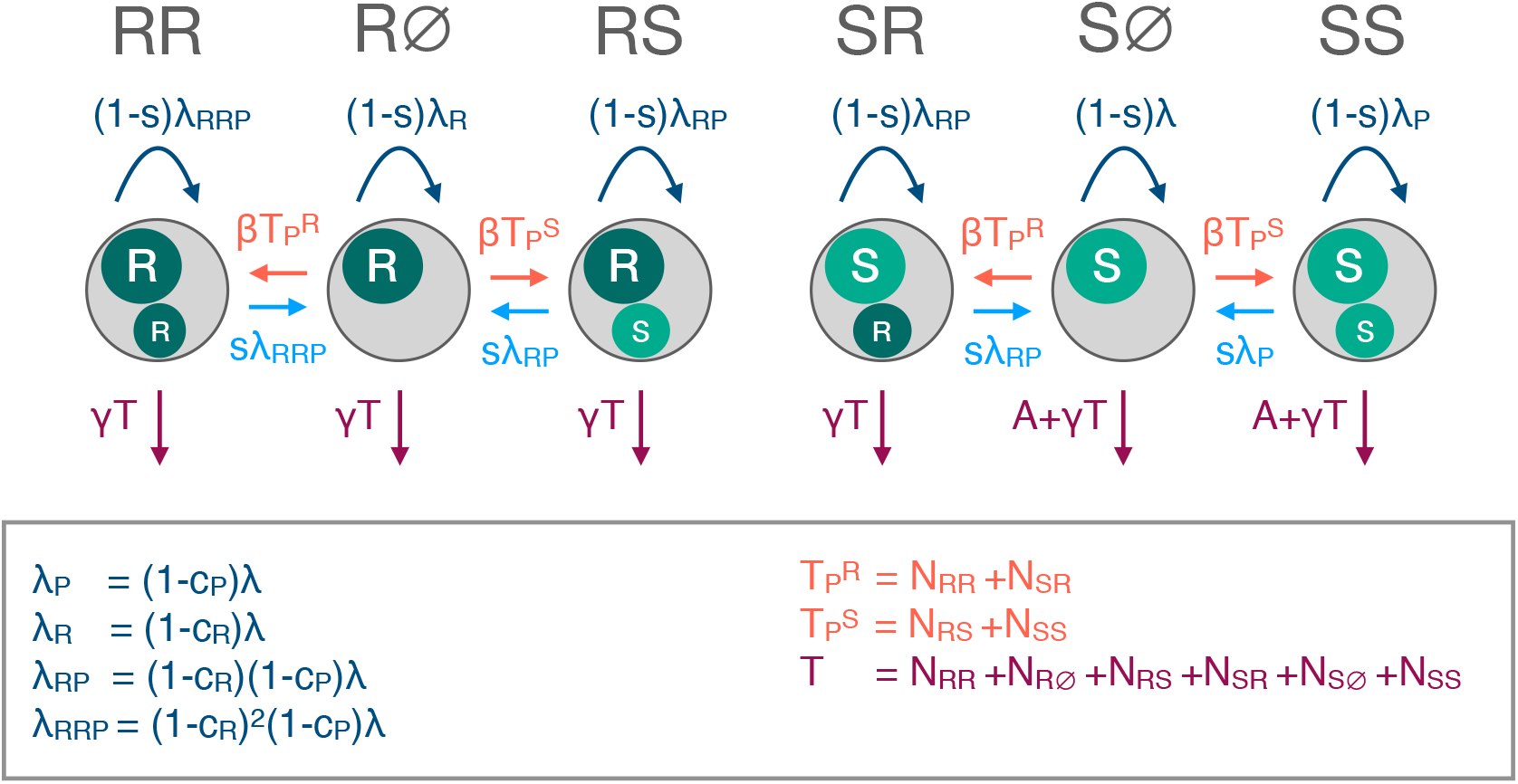
Schematic of the modelled dynamics (Equations 1). Each gray circle represents a cell type, with the interior circles representing the chromosome (large) and plasmid (small), with *R* denoting resistance and *S* sensitivity. The arrows indicate modelled processes: cell replication (dark blue), death (purple), plasmid transmission (orange) and segregation loss (light blue). The labels indicate the rate at which these processes occur. *N* indicates the density of a cell type, so *T* is the total cell density (*T* = *N*_*S*∅_ + *N*_*SS*_ + *N*_*SR*_ + *N*_*R*∅_ + *N*_*RS*_ + *N*_*RR*_), 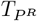 is the total density of cells with a resistant plasmid 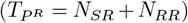, and 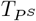 is the total density of cells with a sensitive plasmid 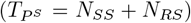. *λ* is the replication rate; *γ* the density dependent death rate; *A* the antibiotic-associated death rate; *β* the plasmid transmission rate; *s* the probability of segregation loss; *c*_*P*_ the cost of plasmid carriage and *c*_*R*_ the cost of carrying the resistance gene.

The arising dynamics are described by the following equations:

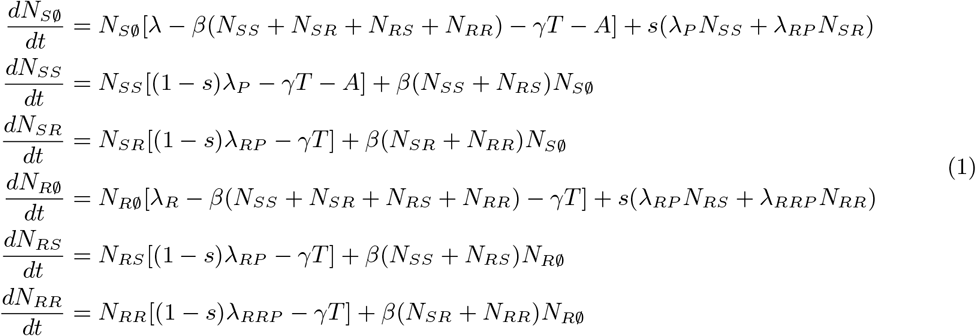

With:

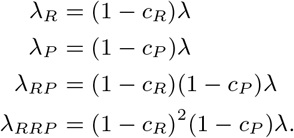

The model parameters are summarised in Table 1. We expect parameter values (i.e. rates and costs) to differ considerably depending on, for example, bacterial species, type of plasmid, antibiotic, and environment. Our aim is to understand the behaviour of a generalised system qualitatively, rather than make quantitative predictions about a specific system. We therefore explore a wide range of parameter values (SI Section 1) rather than choosing parameters to reflect a particular system.

**Table 1:** List of parameters, with their dimensions, the value used in the main text, ranges explored in SI Section 1 and effect on evolutionary outcome. More specifically, the fifth column indicates whether the region of bi-stability (where resistance can be either plasmid-borne or chromosomal) occurs at high or low parameter values compared to the region where only chromosomal resistance is evolutionarily stable (see SI Section 1). The parameter units are arbitrary. The main text values are chosen to best illustrate the range of evolutionary stable outcomes.

| Parameter | Definition | Dimensions | Main text value<br>(SI range) | Bi-stability<br>when |
| --- | --- | --- | --- | --- |
| $\lambda$ | Replication rate | time<sup>-1</sup> | 1 (0.5, 2) | high |
| $\gamma$ | Death rate | volume cells<sup>-1</sup> time<sup>-1</sup> | 1 (0.5, 2) | low |
| $A$ | Antibiotic-associated<br>death rate | time<sup>-1</sup> | 1 (0, 2) | low |
| $c_R$ | Cost of antibiotic<br>resistance | dimensionless | 0.05 (0, 0.5) | high |
| $c_P$ | Cost of plasmid<br>carriage | dimensionless | 0.075 (0, 0.5) | low |
| $\beta$ | Plasmid transmission<br>rate | volume cells<sup>-1</sup> time<sup>-1</sup> | 0.2 (0, 0.25) | high |
| $s$ | Segregation loss | dimensionless | 0.005 (0, 0.1) | low |

### Evolutionary stability of plasmid-borne and chromosomal resistance

We are interested in the evolutionary stability of chromosomal and plasmid-borne resistance, i.e. whether established chromosomal resistance can be displaced by plasmid-borne resistance and vice-versa. We determine parameter regions in which each type of resistance is stable (Figure 2 and SI Figures 1 and 2) using linear stability analysis (see Methods). Under conditions selecting for resistance, we observe three behaviours: evolutionary stability of chromosomal – but not plasmid-borne – resistance; evolutionary stability of plasmid-borne – but not chromosomal – resistance; and evolutionary stability of both forms of resistance. In this third region, resistance occurs on either the plasmid or on the chromosome, but not both: having both chromosomal and plasmid-borne resistance increases the cost of resistance whilst providing no added benefit.

**Figure 2:**
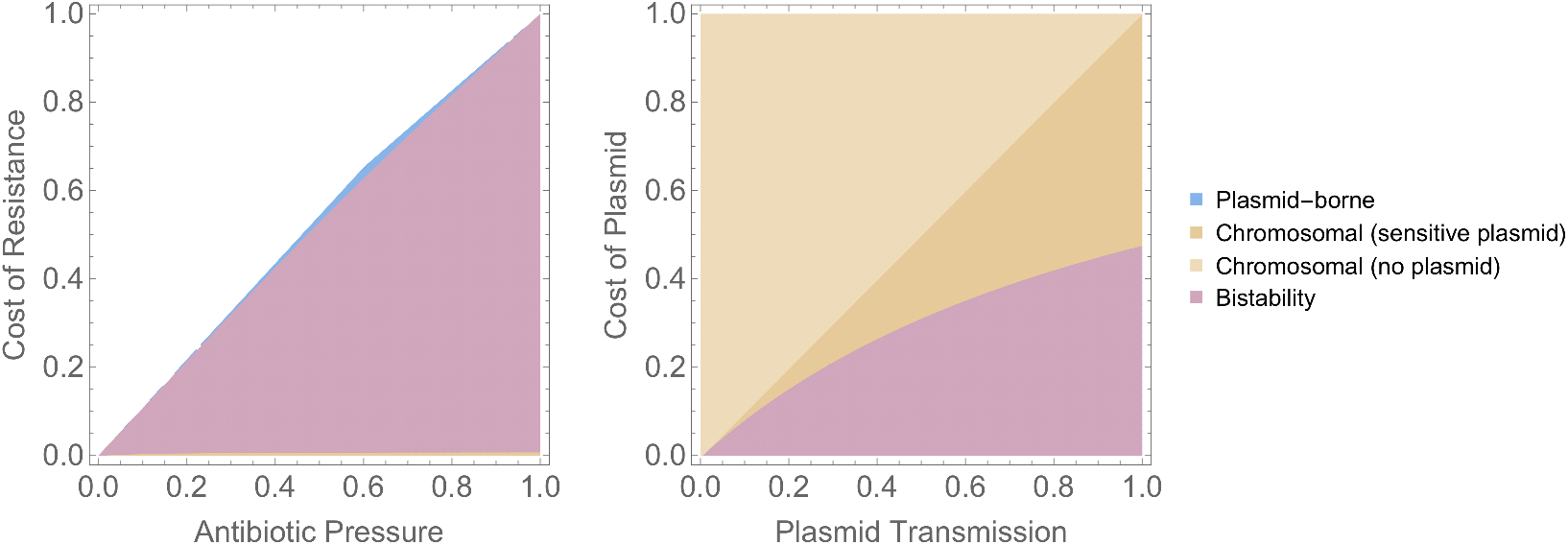
Evolutionary stability of plasmid-borne and chromosomal resistance. The colours indicate which form of resistance is evolutionarily stable: only chromosomal resistance (orange); only plasmid-borne resistance (blue); or either (purple). When resistance is chromosomal, the sensitive plasmid can either be present or absent from the population (dark vs light orange). In the white space in the left-hand panel, neither form of resistance is stable (the population is antibiotic-sensitive). Parameter values are: *λ* = 1, *γ* = 1, *s* = 0.005. For left-hand panel *c*_*P*_ = 0.075 and *β* = 0.2. For right-hand panel *A* = 1 and *c*_*R*_ = 0.05. SI Figures 1 and 2 show results for more parametrisations. Note that in the parameter space where resistance is beneficial, it can either be essential (*A* ≥ *λ*: antibiotic susceptible cells are not viable, even in the absence of competition from resistant cells) or non-essential. This distinction does not impact our results (SI Figures 1 and 2).

#### Chromosomal resistance only

when only chromosomal resistance is evolutionarily stable, resistance genes will always end up on the chromosome over an evolutionary time-scale. The plasmid will either be sensitive, or absent from the population. In general terms (Table 1 and SI Figures 1 and 2), chromosomal resistance is the only evolutionarily stable outcome when the benefit from resistance is high (high antibiotic-associated mortality, low cost of resistance); when the fitness of the plasmid is low (low plasmid transmission rate, high segregation loss, high plasmid cost) and when overall cell density is low (high death rate, low replication rate).

#### Plasmid-borne resistance only

when plasmid borne-resistance is evolutionary stable, resistance genes will always end up on the plasmid. Note that in this region, chromosomal resistance is not stable at all (SI Figure 12): it represents a region in which resistance genes can only persist when horizontally transferred [29]. This outcome only arises under very specific conditions (small parameter space, when resistance yields only a minor fitness benefit, Figure 2) and its presence is sensitive to model structure (for example, how antibiotic effect is modelled, see SI Figure 7). We therefore do not consider this an ecologically plausible explanation for why resistance genes are on plasmids.

#### Bi-stability

when both equilibria are evolutionarily stable, resistance can be either chromosomal or plasmid-borne depending on initial conditions. Once one form of resistance has established, it can no longer be displaced by the other.

To further investigate the dependence on initial conditions, we simulate the system numerically, starting at different initial cell densities (see Methods). We consider a scenario with an initial population consisting of resistant cells and sensitive cells (Figure 3). We vary i) the initial frequency of the sensitive plasmid in the sensitive population; ii) the initial frequency of chromosomal vs plasmid-borne resistance in the resistant population; and iii) whether the chromosomally resistant cells carry the sensitive plasmid. The results of these simulations (Figure 3 and SI Figures 3 and 4) provide insight into the evolutionary pressures that determine the location of resistance genes in three ways.

**Figure 3:**
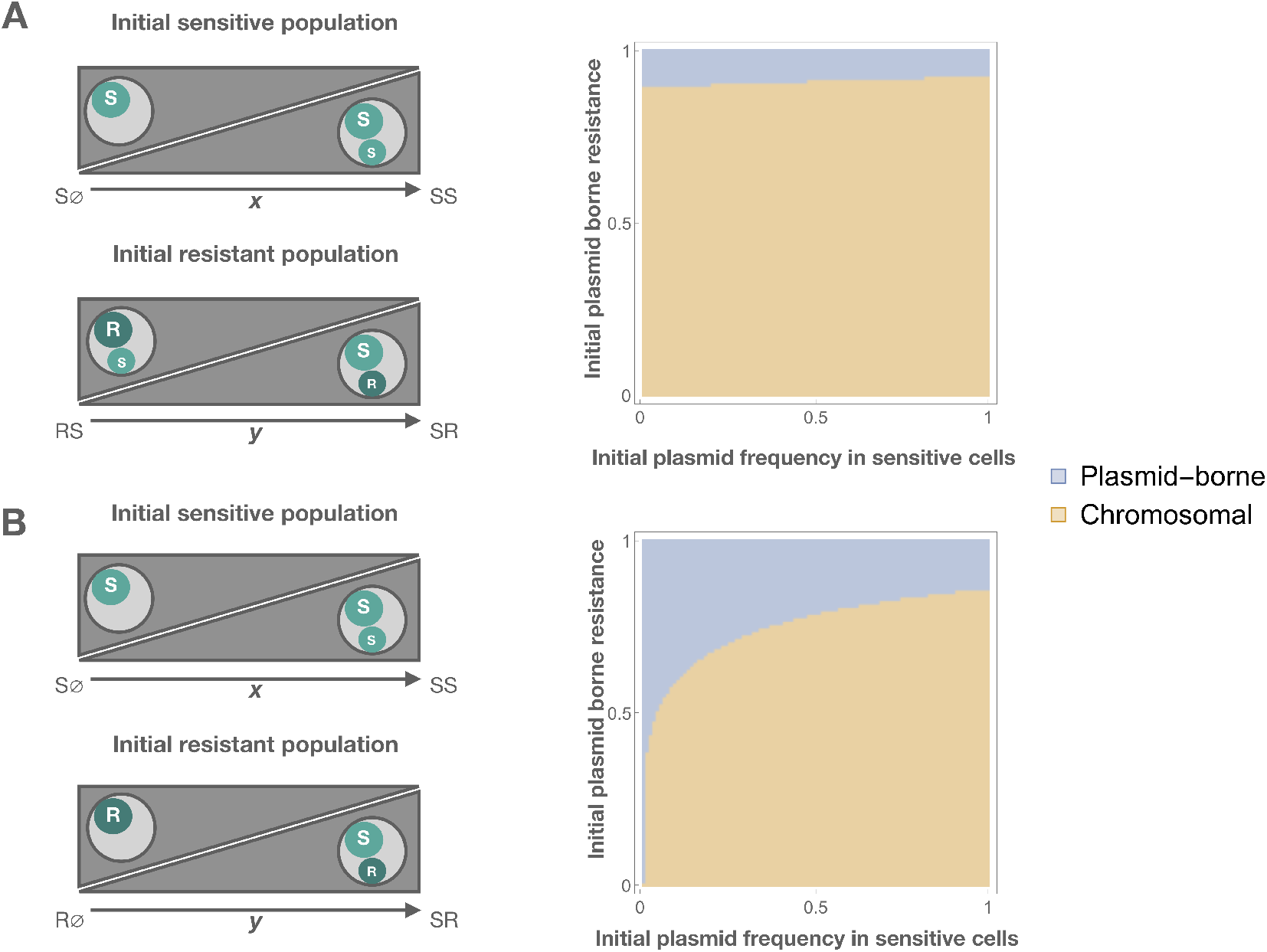
The effect of initial conditions on the equilibrium location of the resistance gene, showing evolutionary outcome depends on both the initial frequencies of plasmid-borne and chromosomal resistance and the initial frequency of the sensitive plasmid. The left-hand panels illustrate variation in the initial conditions. The right hand panels illustrate whether plasmid-borne (blue) or chromosomal resistance (orange) is observed at equilibrium. The x-axis indicates the frequency of the sensitive plasmid in the initial sensitive population *N*_*SS*_*/*(*N*_*S*∅_ + *N*_*SS*_). The y-axis indicates the frequency of the plasmid-borne resistance in the initial resistant population *N*_*SR*_*/*(*N*_*SR*_ + *N*_*RS*_) for panel A, *N*_*SR*_*/*(*N*_*SR*_ + *N*_*R*∅_) for panel B. Plasmid-borne resistance is a more typical outcome in panel B than A because of the presence of the sensitive plasmid in the initial chromosomally resistant population in panel A. The total densities of the initial sensitive and resistant populations are both 1. (Varying the initial ratio of resistance to sensitivity does not affect qualitative results - SI Figure 3). Parameter values are: *λ* = 1, *γ* = 1, *s* = 0.005, *c*_*P*_ = 0.075, *β* = 0.2, *A* = 1 and *c*_*R*_ = 0.05.

Firstly, the presence of positive frequency-dependent selection: plasmid-borne resistance is a more typical outcome when the initial frequency of plasmid-borne resistance is high compared to the frequency of chromosomal resistance. Similarly, a high initial frequency of chromosomal resistance, compared to plasmid-borne resistance, leads to chromosomal resistance as the evolutionary outcome. The fitness of one type of resistance is therefore positively correlated with its frequency. This frequency-dependence arises because dually resistant cells are less fit than cells with either form of single resistance: dual resistance incurs an additional fitness cost but provides no additional fitness benefit. The higher the frequency of chromosomal resistance, the higher the probability that a resistant plasmid will infect a chromosomally resistant (rather than chromosomally sensitive) cell. This disadvantages the resistant plasmid. Similarly, the higher the frequency of the resistant plasmid, the higher the probability that a chromosomally resistant cell will be infected with the resistant plasmid. This disadvantages the resistant chromosome. Thus, the more common the resistance form, the greater its fitness compared to the other form.

Secondly, the evolutionary outcome also depends on the frequency of the sensitive plasmid. Plasmid-borne resistance benefits from the sensitive plasmid being rare: plasmid-borne resistance is a more typical outcome when the initial chromosomally resistant population does not carry the sensitive plasmid and when the frequency of the plasmid in the sensitive population is low. This is because a low initial frequency of the sensitive plasmid means that plasmid-borne resistance can spread both vertically (cell replication) and horizontally (plasmid transmission), allowing it to increase in frequency more rapidly than chromosomal resistance.

Thirdly, overall, chromosomal resistance is a more typical outcome in these simulations than plasmid-borne resistance. This is because plasmid-borne resistance, unlike chromosomal resistance, is subject to segregation loss: it is not always inherited during cell replication. Indeed, increasing the probability of segregation loss favours chromosomal resistance (SI Figure 4).

#### Robustness of results

We test the robustness of these results to a number of assumptions about model structure (see Methods and SI Section 2). The general result is that the presence of bi-stability is robust, although the size of the region of bi-stability can change. The only crucial assumption for positive frequency-dependence is that dual resistance is less beneficial than single resistance (SI Figure 5 and 6). In other words, eliminating the additional cost from dual resistance, or increasing the benefit of dual resistance so much that it outweighs this additional cost, abolishes the region of bi-stability. Under these circumstances, dual resistance dominates (i.e. the population will consist of a resistant plasmid circulating in a chromosomally resistant population).

The results of two sensitivity analyses in particular are worth highlighting. Firstly, our results are robust to inclusion of gene flow between the plasmid and chromosome (e.g. transposition of the resistance gene). Gene flow allows the otherwise excluded form of resistance to persist at low frequency (analogous to mutation-selection balance), and increases the range of initial conditions leading to chromosomal resistance (SI Figure 8). However, these effects only become substantial for unrealistically high transposition rates (SI Section 2.4) [30]. Secondly, the presence of bi-stability is robust to modelling fluctuating, instead of constant, antibiotic pressure. Depending on its period, fluctuation can favour plasmid-borne resistance, increasing the size of the parameter space in which only plasmid-borne resistance is evolutionarily stable (SI Figure 10).

#### Relationship to previous modelling results

Next, we revisit some previous modelling results. As discussed in the introduction, previous modelling predicts that locally beneficial traits will be plasmid-borne rather than chromosomal, thus providing a complementary hypothesis for why certain genes reside on plasmids [16]. However, the model from which this prediction is derived assumes absence of the plasmid outside the local niche. We therefore ask whether local adaptation favours plasmid-borne resistance if the sensitive plasmid can persist outside the local niche. We modify our model to include an influx of sensitive cells, and vary the frequency of the sensitive plasmid in these incoming cells (SI Section 3). This corresponds to a scenario in which resistance is locally beneficial in the modelled environment, but not selected for elsewhere. As shown in Figure 4, an influx of sensitive cells without the sensitive plasmid does indeed favour plasmid-borne resistance, as suggested previously [16]. However, an influx of sensitive cells with the sensitive plasmid favours chromosomal resistance. The strength of this effect depends on the rate of influx of sensitive cells. Thus, local adaptation only favours plasmid-borne resistance if the frequency of the sensitive plasmid is low outside the local niche.

**Figure 4:**
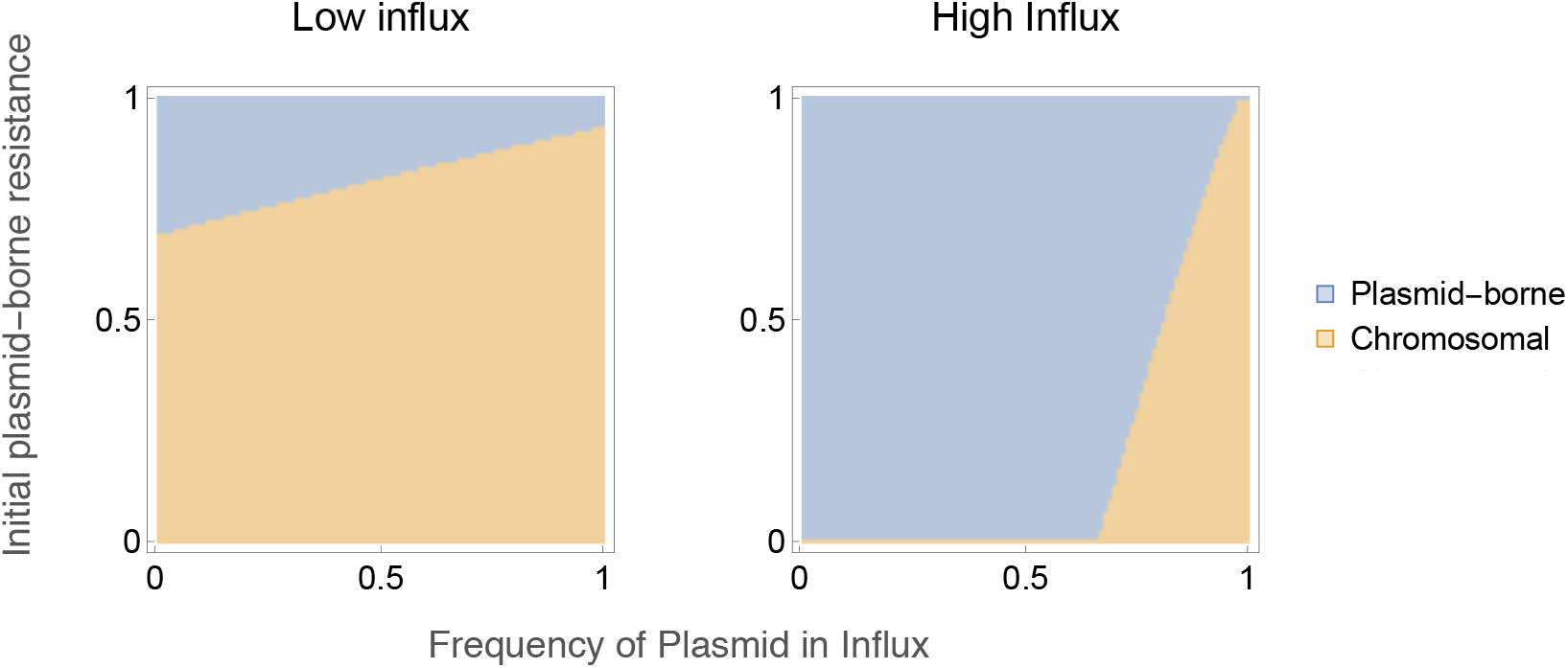
The location (chromosomal or plasmid-borne) of a locally beneficial resistance gene depends on the presence of the sensitive plasmid in immigrant cells. The initial population is fully resistant (with chromosomally resistant cells carrying the sensitive plasmid, corresponding to panel A in Figure 3), with the y-axis indicating the frequency of plasmid-borne resistance in this initial population *N*_*SR*_*/*(*N*_*SR*_ + *N*_*RS*_). The x-axis indicates the frequency of the sensitive plasmid in the immigrant cells. The presence of the plasmid in these immigrant cells favours chromosomal resistance. The high influx rate is *μ* = 10^−1^, the low influx rate is *μ* = 10^−2^. Other parameters values are: *λ* = 1, *γ* = 1, *s* = 0.005, *c*_*P*_ = 0.075, *β* = 0.2, *A* = 1 and *c*_*R*_ = 0.05.

Secondly, we revisit results relating to plasmid persistence. Previous modelling work has suggested that if plasmid fitness is too low for plasmids to persist as pure parasites (i.e. without carrying genes beneficial to the host cell), beneficial genes will always locate on the chromosome rather than plasmid (in absence of local adaptation) [16]. Thus, the persistence of low transmissibility plasmids is a paradox: they cannot be maintained without beneficial genes, but beneficial genes cannot be maintained on these plasmids [16].

We test this prediction in our model (as detailed in SI Section 4) by comparing the parameter space in which plasmid-borne resistance is evolutionarily stable (i.e. resistance genes can locate onto the plasmid even in the presence of competition from chromosomal resistance) with the parameter space in which a parasitic plasmid can persist (i.e. a sensitive plasmid can persist in a chromosomally sensitive population). We find that previous results do not hold for the model structure presented here: resistance genes can locate onto the plasmid instead of the chromosome even if the plasmid transmissibility is too low for the plasmid to persist as a parasite (SI Figure 11). This implies that in theory, it is possible for there to be low transmissibility plasmids which persist purely because of the advantage they provide host cells. It is worth noting, however, that the parameter space in which this occurs is small (SI Figure 11).

#### Rate of acquisition determines resistance gene location

Thus far, our results show that for moderately beneficial genes (i.e. those in the bi-stable parameter region), the presence of positive frequency-dependent selection means that plasmid-borne resistance can be evolutionarily stable despite segregation loss. This frequency-dependent selection is not, in itself, a sufficient explanation for why resistance genes are plasmid-borne. However, it does suggest that whichever form of resistance (plasmid-borne or chromosomal) is acquired first is likely to establish in the population: if the first form of resistance has time to increase in density prior to the acquisition of the other form, its greater frequency will give it a fitness advantage. The first resistance type need not have reached fixation in order to preclude invasion by the other: the frequency-dependent advantage is sufficiently strong even at low overall resistance frequencies (SI Figure 13). Therefore, when the rate of resistance acquisition is low compared to the rate of increase in resistance frequency once acquired, the first form of resistance will persist.

Thus, the presence of resistance genes on plasmids could be explained by the acquisition rate of plasmid-borne resistance being higher than the acquisition rate of chromosomal resistance. Indeed, rates of conjugative plasmid transfer are generally higher than rates of chromosomal horizontal gene transfer (one estimate, based on comparison of experimental measures, suggests of the order of 10^7^ higher, though this is probably highly context dependent [31]). Furthermore, for a number of bacterial species, the primary mechanism of resistance gene acquisition is indeed thought to be inter-species transfer of resistance-bearing plasmids [32, 33].

To formalise this idea, we develop a simple model of resistance acquisition in multiple species (Figure 5 and Methods). We model *n* species; a resistance gene which is beneficial in all species; and a plasmid that can be transferred between and persist in all species (either because it has a broad host range or because its range can be shifted or expanded following transfer [34]). Resistance can be either plasmid-borne or chromosomal. Once a species acquires one form of resistance, this form of resistance becomes established and can no longer be replaced (due to positive frequency-dependent selection). We assume resistance genes only emerge *de novo* on the chromosome (at rate *m*). The gene can spread through inter-species horizontal transfer of chromosomal resistance (e.g. transformation) (at rate *c*), or inter-species transfer of resistance plasmids (at rate *p*). We assume the gene can move between the plasmid and chromosome at low rates, which allows the otherwise excluded form of resistance to persist at low frequency. We do not explicitly model this coexistence, but do model the horizontal transfer of the low frequency form (at rate *t* ∗ *p* for plasmid-borne resistance and rate *t* ∗ *c* for chromosomal resistance, where *t* is the frequency of the low frequency form).

**Figure 5:**
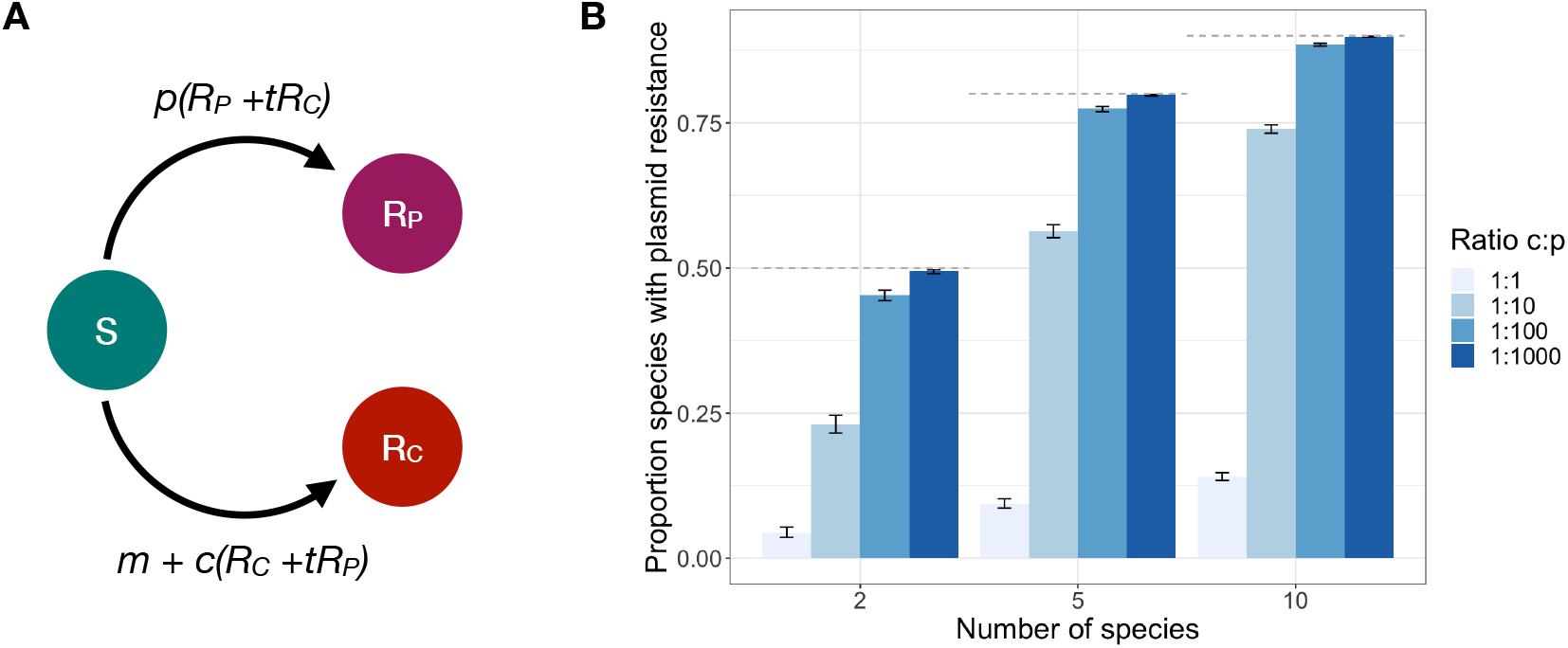
Prevalence of plasmid-borne resistance. Panel A: Representation of the model structure. *S* represents species without the resistance gene; *R*_*P*_ species with the resistance gene on the plasmid; *R*_*C*_ species with the resistance gene on the chromosome. *m* is the rate at which resistance arises through mutation; *p* the rate at which the plasmid-borne resistance is transferred between species; *c* the rate at which chromosomal resistance is transferred between species; and *t* captures gene flow between the plasmid and chromosome. Panel B: The proportion of plasmid-borne resistance depends on the number of simulated species and the ratio of the rate of interspecies transfer of the chromosomal (*c*) and plasmid-borne gene (*p*). The horizontal dashed lines show the maximum proportion of plasmid resistance, given that resistance must first emerge on the chromosome ((*n* − 1)*/n*). The error bars represent 95% confidence intervals, based on 1000 realisations. Parameters were: *m* = 10^−6^, *c* = 10^−5^ and *t* = 10^−1^. Results for alternative parameterisations are shown in SI Figure 14.

We simulate this system stochastically (see Methods), starting from no species having the resistance gene. Figure 5 shows the proportion of species with plasmid-borne resistance once the gene has spread to all species. As expected, the proportion of species with plasmid-borne resistance increases with the rate of inter-species plasmid transfer. In addition, this proportion also increases with the number of modelled species. This effect arises for two reasons. Firstly, the initial *de novo* appearance of the gene must be on the chromosome. Thus, for example, when only two species are modelled, plasmid-borne resistance can only occur in 1 out of 2 species. Secondly, the impact of the rate of inter-species transfer increases with the number of potential donor species. These results are robust to different parametrisation (SI Figure 14).

## Discussion

To understand why certain gene functions, including antibiotic resistance, are found on plasmids, we develop a model of the competition between resistant and sensitive plasmids and resistant and sensitive chromosomes. Our key finding is that this model gives rise to positive frequency-dependent selection on gene location. This positive frequency-dependence arises when carrying both chromosomal and plasmid-borne resistance is disadvantageous (i.e. the cost of the second copy out-weights any additional reduction in antibiotic-associated death). This disadvantage acts as a barrier to a low frequency form of resistance invading the population. Although the model was formulated in terms of the antibiotic resistance, we expect this central result to generalise to any gene where cells with both chromosomal and plasmid-borne versions are at a disadvantage compared to cells with a single version.

The consequence of this frequency-dependence is that for some parameter ranges genes can be maintained on plasmids, despite segregation loss, if they start with a frequency advantage. The key conditions are that the gene is only moderately beneficial – highly beneficial genes, whether strictly ‘essential’ or not, are always chromosomal, because segregation loss is too detrimental – and that the fitness of the plasmid is not too low. Under these conditions, gene location (i.e. chromosomal or plasmid-borne) depends on which form is acquired first. Using a simple stochastic model of resistance acquisition and transfer across multiple species, we show that the probability of finding genes on plasmids increases with the rate of inter-species plasmid transfer and with the number of species between which the gene and plasmid can be shared.

We also revisit the previously proposed idea that plasmids code for locally beneficial genes, and show that local adaptation does not explain plasmid-borne resistance when the sensitive plasmid is present outside the local niche. Yet, many plasmid-borne traits, such as antibiotic resistance, but also heavy metal tolerance [8] or metabolism of rare substances [5], seem to fit the description of local or intermittent usefulness. Our results raise the possibility that these genes are on plasmids not because they code for local adaptation as such, but because their local usefulness means they are, on average, only moderately beneficial.

All together, the hypothesis that emerges from these results is that plasmid-borne genes are i) moderately beneficial, possibly due to heterogeneous selection pressure; ii) functional across a large number of species; and iii) rarely acquired through chromosomal mutation; and that resistance genes are often found on plasmids because these genes commonly fulfill these criteria.

Our model considers a single antibiotic and resistance gene. Resistance plasmids often carry multiple genes encoding resistance to different antibiotics [35]. As higher plasmid fitness favours plasmid-borne resistance in our model, we might expect resistance genes to accumulate on highly transmissible plasmids. Furthermore, as acquisition of one beneficial gene increases plasmid fitness, such acquisition might allow the plasmid to aggregate further genes. However, it should also be noted that there are a number of other evolutionary mechanisms, in particularly correlated selection pressure for resistance against different antibiotics, which contribute to the co-occurrence of resistance genes [36, 37].

Throughout the work, we have made the assumption that the effect of the gene of interest (e.g. for antibiotic resistance genes, their cost and effectiveness against antibiotic-associated mortality) is the same on the chromosome and the plasmid: when this assumption does not hold, location can be trivially explained by the fitness difference. This assumption refers specifically to the effects of the same gene on the plasmid and chromosome when the gene is first introduced. (Note that a meta-analysis of chromosomally acquired and plasmid-acquired antibiotic resistance found the former to be costlier on average [38], but the analysis did not compare the cost of the *same* gene on the plasmid and chromosome.) It is relevant to consider this equal effect assumption in the light of both plasmid copy number and compensatory evolution.

Firstly, the equal effects assumption would not hold for high copy number plasmids if gene expression levels increase with copy number: increased expression may lead to both higher cost (higher metabolic burden) and higher effectiveness. However, the extent to which gene expression increases with plasmid copy number is unclear because it depends on how tightly gene expression is regulated, which varies between plasmid genes [39]. Indeed, while there are some examples of phenotypic resistance increasing with copy number (e.g amoxicillin-clavulanate resistance in *Escherichia coli* [40]), evidence from *Klebsiella pneumoniae* suggests that whether this effect is observed depends on both the antibiotic-gene combination and the genetic background [41].

Secondly, compensatory evolution may eventually lead to genes having a different cost on the plasmid and chromosome, for example, through regulation of gene expression [26, 39]. However, we would not expect compensatory effects to be present when the gene is first introduced into the species. Indeed, compensatory evolution might subsequently reinforce the priority effect we observe, by giving the first form of resistance more time to acquire compensatory mutations [38].

The prediction of bi-stability is experimentally testable: this requires demonstrating that there are conditions under which resistance genes can be either chromosomal or plasmid-borne, and that neither form of resistance can invade a population in which the other form of resistance is established. More specifically, our model predicts bi-stability when plasmid transmissibility is high and the benefit of the gene low: for lower transmissibility plasmids and a higher benefit from resistance (e.g. higher antibiotic concentration), we would expect chromosomal resistance to always invade.

Although we are not aware of studies directly testing these effects, the model predictions are consistent with existing experimental findings. Firstly, previous experimental work has demonstrated that whether chromosomal resistance invades a population with plasmid-borne resistance depends on plasmid transmissibility, with low plasmid transmissibility allowing invasion [42, 43]. This is consistent with our model’s predictions. However, demonstrating the presence of bi-stability would require also showing that plasmid-borne resistance cannot invade chromosomal resistance even when plasmid transmissibility is high.

Secondly, in a study of compensatory evolution to alleviate the fitness cost of a plasmid carrying mercury resistance, Harrison et al. found that the resistance gene frequently transitioned to the chromosome [26]. These chromosomally resistant lineages lost the plasmid, but were generally not able to increase in frequency. The authors attributed the effect to compensatory evolution alleviating the cost of the plasmid. However, unless the cost of the plasmid was fully alleviated, the chromosomally resistant and plasmid-free lineage would still be fitter than the plasmid-bearing lineage and thus expected to invade. Yet, if the cost of the plasmid was fully alleviated, the chromosomally resistant lineage would not necessarily lose the plasmid. Therefore, the inability of these chromosomally resistant cells to invade is compatible with both compensatory adaptation and frequency-dependent selection.

Rodrìguez-Beltrán et al [44] have recently shown that polyploidy (i.e. gene copies being present on both mobile element and chromosome or, for multi-copy plasmids, on multiple plasmids) plays an important, and thus far neglected, role in mobile genetic element evolution: polyploidy masks the effect of recessive mutations on mobile genetic elements (‘genetic dominance’), and thus mobile genetic elements are primarily associated with dominant mutations. Our results can also be thought of in terms of polyploidy: positive frequency dependence arises because polyploidy has different effects on the cost and effectiveness of the gene. Our results are therefore another example of the emerging importance of polyploidy effects in understanding the gene content of plasmids.

Finally, our results provide a new perspective on the extent to which the long-term fate of resistance genes depends on stochastic gene acquisition events. Recent analysis of resistance dynamics in *Streptococcus pneumoniae* has suggested that the rate at which lineages acquire resistance genes is not a major determinant of their resistance frequencies. This supports the view that resistance evolution is a deterministic outcome of selection pressures [45]. Our results provide a different perspective: in the context of the location of resistance genes, the priority effect arising from positive frequency-dependent selection means that the timing of gene acquisition events sets the population onto an evolutionary path from which it cannot subsequently deviate. Thus, in this context, eventual evolutionary outcomes are fundamentally stochastic.

## Methods

### Computation of equilibria and stability analysis

We consider the dynamics of a population consisting of cells which can be either chromosomally resistant or chromosomally sensitive, with no plasmid, a resistant plasmid, or a sensitive plasmid, giving rise to a model with six cell types (Equations 1 and Figure 1). Our aim is to study the evolutionary stability of plasmid-borne and chromosomal resistance by finding the equilibria of this system and determining their stability. We were unable to solve for the equilibria of the full six species system. However, we can use insights into dynamics of the system to simplify the problem. Firstly, we do not expect coexistence of the sensitive and resistant chromosome, nor of the sensitive and resistant plasmid: in each case, the sensitive and resistant variant are competing for the same resource and thus, without a coexistence promoting mechanism, we expect to see competitive exclusion [46, 47, 48]. Secondly, we do not expect presence of cells with both plasmid-borne and chromosomal resistance at equilibrium (*RR*): because dual resistance carries a dual cost, but not a dual benefit, these cells will always be inferior to cell with single resistance (*R*∅, *RS* or *SR*).

This allows us to narrow the possible equilibria in which resistance is present to three: presence of chromosomal resistance, either in presence or absence of the sensitive plasmid (*N*_*R*∅_ *>* 0, *N*_*RS*_ *>* 0, all other cell types 0; or *N*_*R*∅_ *>* 0, all other cell types 0), and plasmid-borne resistance (*N*_*S*∅_ *>* 0, *N*_*SR*_ *>* 0, all other cell types 0). For each equilibrium of interest, therefore, we reduce the full model by setting the relevant subset of the variables to zero and compute the steady state of this reduced system. We then check the stability of this steady state in the *full model* by computing the eigenvalues of the full model’s Jacobian evaluated at this equilibrium point. The parameter region in which all six eigenvalues are negative is the region in which the equilibrium is evolutionarily stable.

These calculations were implemented in Wolfram Mathematica 12 [49]. The code is available as a Supporting File. Note that our reasoning for narrowing down the equilibrium of interest is verbal rather than a mathematical: we can therefore not rule out the possibility we have overlooked a relevant equilibrium. However, such equilibria are never seen in any of our numerical simulations.

### Numerical simulations of dynamics

For the analyses examining the dependence of resistance on initial conditions, we simulate the system numerically until equilibrium is reached (for 10^7^ timesteps unless otherwise indicated). To avoid non-zero cell densities arising from numerical errors, final states are rounded to 10^−10^. These simulations were implemented in Wolfram Mathematica 12 [49]. The code is available as a Supporting File.

### Sensitivity analyses

We perform sensitivity analyses to test the robustness of our results to a number of changes in model structure: modifying the effect of dual resistance (decreasing the cost, increasing the benefit); modelling segregation loss independently from replication; modelling the antibiotic as slowing growth rate rather than increasing death rate; modelling fitness costs as increased death rate rather than decreased growth rate; allowing gene flow between the plasmid and chromosome; and relaxing the assumption that carriage of one plasmid completely excludes the other. Full details are provided in SI Section 2.

### Multi-species model

For the multi-species model of resistance acquisition, we model *n* species which can be in one of three states: without resistance (*S*), with chromosomal resistance (*R*_*C*_) or with plasmid-borne resistance (*R*_*P*_). Once species have reached either state *R*_*C*_ or *R*_*P*_, they remain in this state due to frequency-dependent selection favouring the first acquired form of resistance. Species can transition from state *S* to state *R*_*C*_ through *de novo* mutation (at rate *m*) or transfer of the resistance gene from a species with chromosomal resistance (at rate *cR*_*C*_). In addition, we allow for gene flow between the plasmid and chromosome leading to the otherwise excluded form of resistance persisting at low frequency. We do not explicitly model this coexistence, but do model the horizontal transfer of the low frequency form. Thus, species can also transition from state *S* to state *R*_*C*_ through transfer from a species with plasmid-borne resistance (at rate *t* ∗ *c*, where *t* captures the frequency of the low frequency form arising from gene movement between the plasmid and chromosome). Species can transition from state *S* to *R*_*P*_ through inter-species plasmid transfer, either from a species with plasmid-borne resistance (at rate *p*) or chromosomal resistance (at rate *t* ∗ *p*).

We simulate this model stochastically, starting from all species in state *S*, until all species have acquired resistance. If *N*_*R*_*C*__ is the number of species with chromosomal resistance and *N*_*R*_*P*__ is the number of species with plasmid-borne resistance, then, at any given timestep, the probability of any given species transitioning from the *S* state to one of the *R* states is given by:

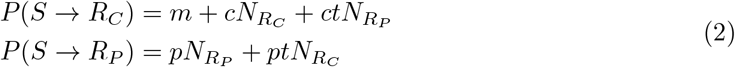

Figure 5 is based on 1000 realisations. The simulation was implemented in R [50] and the code is available as a Supporting File.

## Acknowledgments

We thank Alex Hall for comments on the manuscript. The study was funded by the Swiss National Science foundation (grant number 310030B 176401 and NRP72 SNF grant number 407240-167121). The authors declare that there is no conflict of interest.

## Data availability

The data underlying this article are available in the article and in its online supplementary material.

## Supporting Information

### 1 Parametrisation

#### 1.1 Evolutionary Stability

SI Figures 1 and 2 explore the evolutionary stability of plasmid-borne and chromosomal resistance for a wide range of parameter combinations. Each figure is associated with a set of standard parameter values, with each panel exploring variation around these standard values for one pair of parameters. For SI Figure 1, the standard parameter values are the same as in main text Figure 2: (*λ* = 1, *γ* = 1, *s* = 0.005, *c*_*P*_ = 0.075, *β* = 0.2, *A* = 1 and *c*_*R*_ = 0.05). SI Figure 2 explores behaviour at a lower antibiotic consumption rate (*A* = 0.5), where sensitive cells are viable at the standard value of the replication rate (*λ* = 1). The two figures are very similar, with minor differences for some panels in which the fitness cost of resistance is varied. Thus, the distinction between essential and simply beneficial resistance is not meaningful in our model.

**Figure 1:**
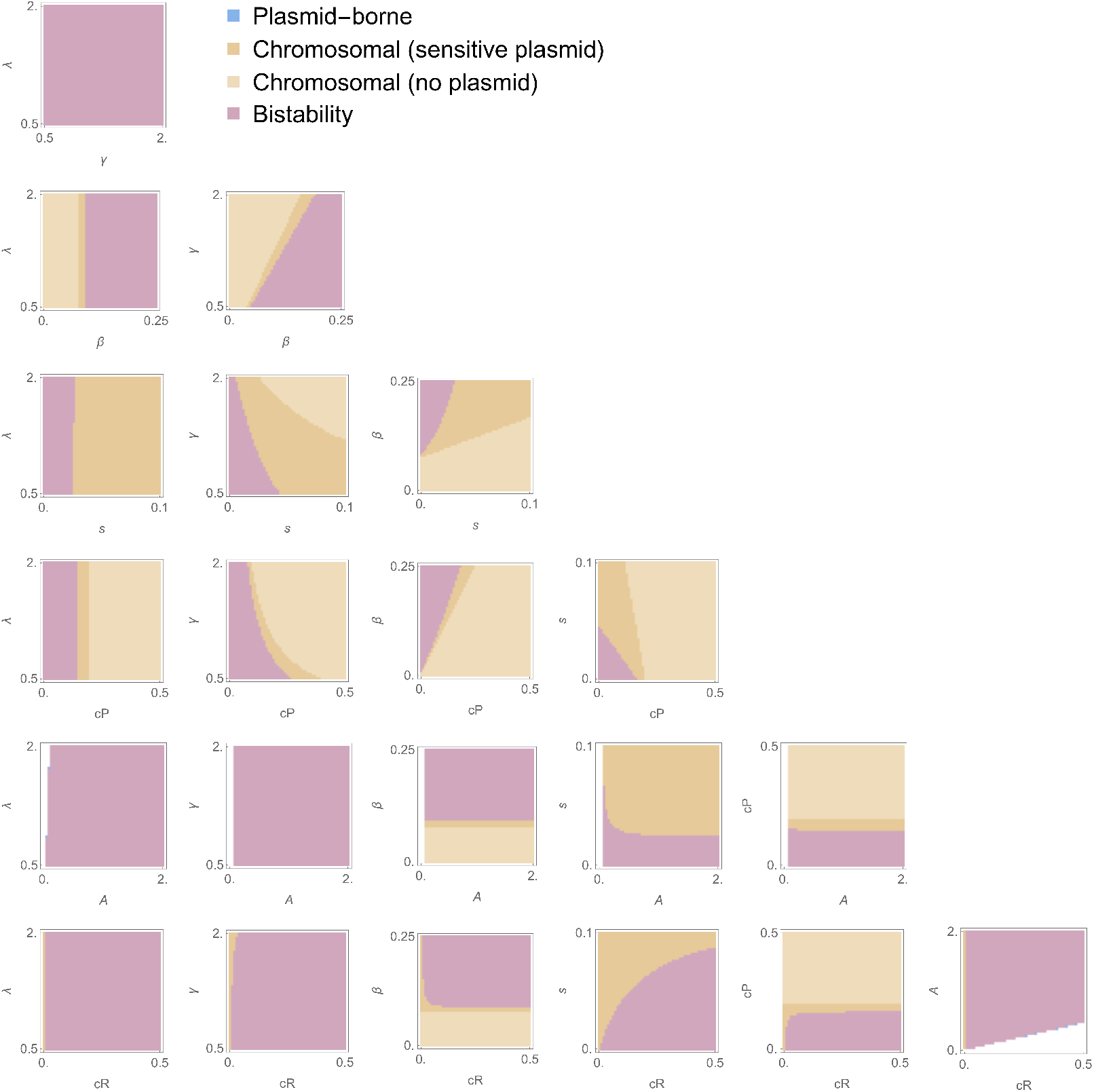
Evolutionary stability of plasmid-borne and chromosomal resistance. Each panel shows parameter regions in which plasmid-borne only (blue); chromosomal resistance (orange); or both (purple) are evolutionarily stable, for varying values of a pair of parameters. For each panel, values for non-varying parameters are the same as in main text: *λ* = 1, *γ* = 1, *s* = 0.005, *c*_*P*_ = 0.075, *β* = 0.2, *A* = 1 and *c*_*R*_ = 0.05.

**Figure 2:**
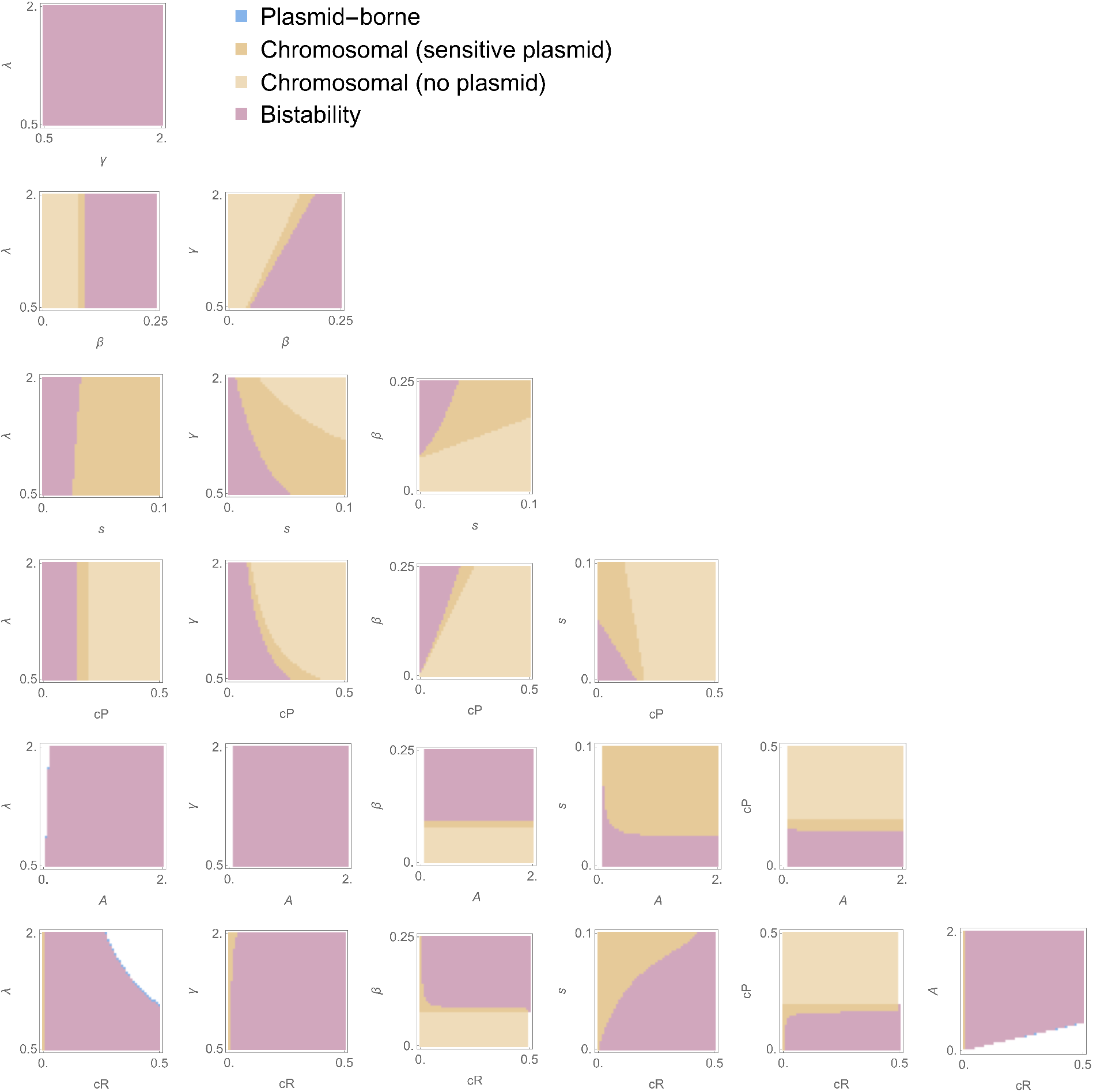
Evolutionary stability of plasmid-borne and chromosomal resistance at lower antibiotic consumption rate (*A* = 0.5), where sensitive cells are viable the standard value of the replication rate (*λ* = 1). Each panel shows parameter regions in which plasmid-borne only (blue); chromosomal resistance (orange); or both (purple) are evolutionarily stable, for varying values of a pair of parameters. For each panel, values for non-varying parameters are the same as in main text, except that antibiotic consumption is lower (and cells without the resistance gene are therefore viable): *λ* = 1, *γ* = 1, *s* = 0.005, *c*_*P*_ = 0.075, *β* = 0.2, *A* = 0.5 and *c*_*R*_ = 0.05.

#### 1.2 Dependence on initial conditions

SI Figures 3 and 4 explore how the relationship between initial conditions and evolutionary outcomes depends on parametrisation. The default parameters are the same as in the main text (*λ* = 1, *γ* = 1, *s* = 0.005, *c*_*P*_ = 0.075, *β* = 0.2, *A* = 1 and *c*_*R*_ = 0.05), with variation indicated in figure labels and legends. The relationship between parameter values and evolutionary outcome is the same as found in the analysis of evolutionary stability, i.e. plasmid-borne resistance is favoured by:

- high replication rate (*λ*), plasmid-transmission rate (*β*), and high cost of resistance (*c*_*R*_)
- low density-dependent death rate (*γ*), segregation loss (*s*), cost of plasmid (*c*_*P*_), and antibiotic associated mortality (*A*).

**Figure 3:**
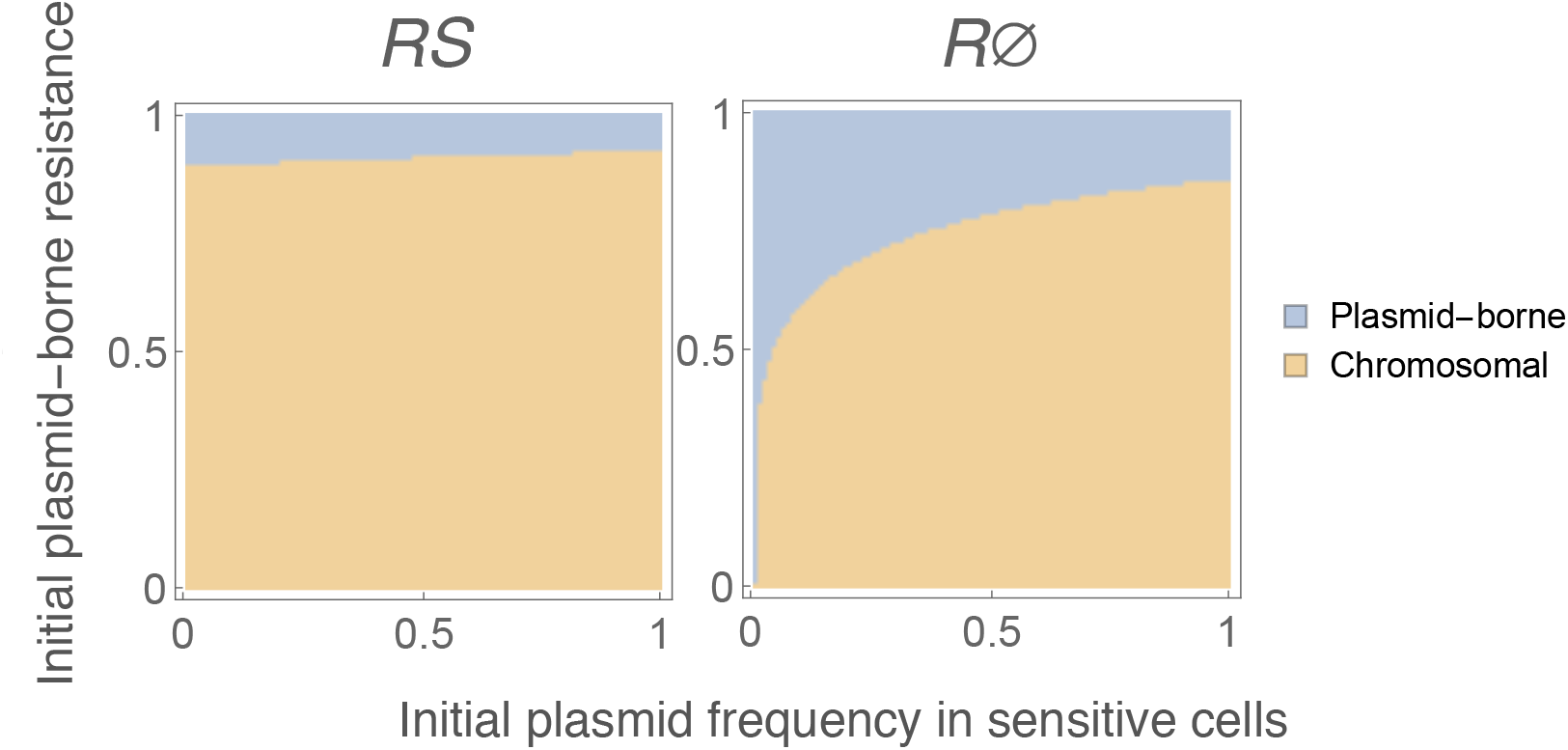
Effect of initial conditions on equilibrium location of resistance gene, with initially low resistance frequency. In main text Figure 3, the initial densities of the sensitive and resistant population are both 1. Here, the initial density of the sensitive population is 1, and the initial density of the resistant population is 0.01. As in main text Figure 3, the panels illustrates whether plasmid-borne (blue) or chromosomal resistance (orange) are observed at equilibrium. The x-axis indicates the frequency of the sensitive plasmid in the initial sensitive population. The y-axis indicates the frequency of the plasmid-borne resistance in the initial resistant population. For the left hand panel, the initial chromosomally resistant population carries the sensitive plasmid (*RS*), for the right hand panel, it does not (*R*∅). Parameter values are: *λ* = 1, *γ* = 1, *s* = 0.005, *c*_*P*_ = 0.075, *β* = 0.2, *A* = 1 and *c*_*R*_ = 0.05.

**Figure 4:**
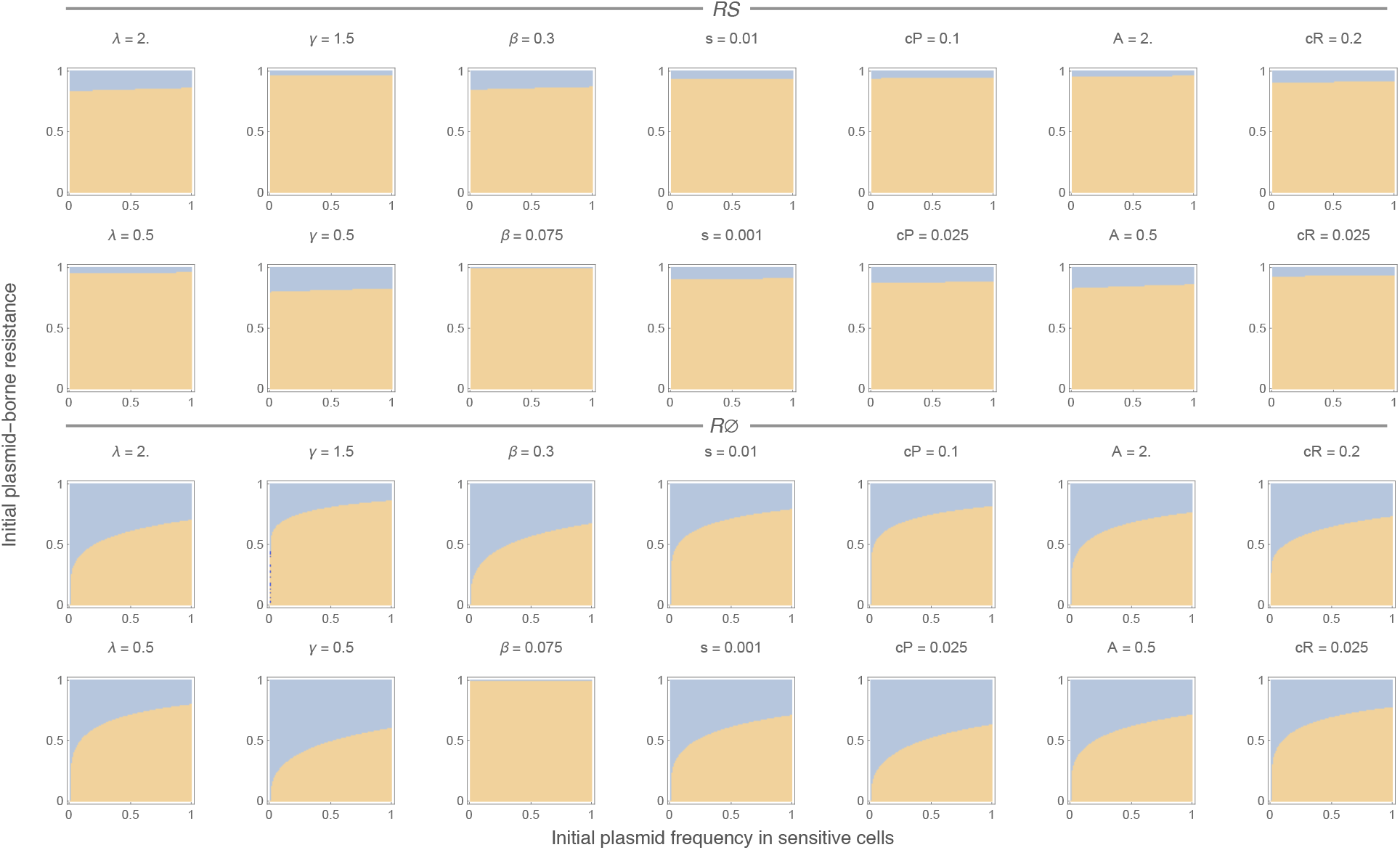
Effect of parametrisation on the relationship between initial conditions and evolutionary outcomes. Each panel depicts the effect of changing one parameter from the default parametrisation (*λ* = 1, *γ* = 1, *s* = 0.005, *c*_*P*_ = 0.075, *β* = 0.2, *A* = 1 and *c*_*R*_ = 0.05). As in main text Figure 3, the panels illustrates whether plasmid-borne (blue) or chromosomal resistance (orange) are observed at equilibrium. The x-axis indicates the frequency of the sensitive plasmid in the initial sensitive population. The y-axis indicates the frequency of the plasmid-borne resistance in the initial resistant population. The total densities of the initial sensitive and resistant populations are both 1. Top rows: chromosomally resistant cells in the initial population carry the sensitive plasmid. Bottom rows: chromosomally resistant cells in the initial population do not carry the sensitive plasmid. For a small number of initial conditions, the numerical simulation of the system failed due to problems with precision (*γ* = 1.5 with initial *R*∅ population, panel in second column, third row). These simulations are indicated in dark blue.

### 2 Sensitivity Analyses

In this section, we test the robustness of our results to assumptions about model structure. SI Table 1 summarises the analyses and their effect on our key results.

**Table 1:** Summary of sensitivity analyses and their effects on model behaviour.

| Sensitivity analysis | Positive frequency-dependent selection (i.e. region of bi-stability) | Other effects |
| --- | --- | --- |
| Decreasing cost of dual resistance | Not observed when dual resistance incurs no additional cost | Favours chromosomal resistance |
| Increasing benefit of dual resistance | Not observed when benefit of dual resistance out-weights its cost | Favours chromosomal resistance |
| Independent segregation loss | Robust | Favours chromosomal resistance |
| Antibiotic effect on growth | Robust | Removes parameter space in which plasmid-borne resistance alone is stable |
| Fitness costs as death rate | Robust | Favours chromosomal resistance; removes parameter space in which plasmid-borne resistance alone is stable |
| Gene flow between plasmid and chromosome | Robust | Allows low level coexistence of both resistance forms; favours chromosomal resistance at high levels of gene flow |
| Imperfect exclusion of plasmid co-infection | Robust | Favours plasmid-borne resistance |
| Fluctuating antibiotic selection | Robust | Favours plasmid-borne resistance under some conditions |

Note that the parametrisation for the figures in this Section (SI Figures 5 to 7) is slightly different from that used the main text: the plasmid transmission rate in the main text is *β* = 0.2, whereas here we use *β* = 0.10. This is to more clearly illustrate that the parameter region at low cost of resistance in which only chromosomal resistance is evolutionarily stable also arises in the main model structure. In other words, that this effect is not a consequence of modifying the model structure. This region is also present when *β* = 0.2 (Main Text Figure 2), but is very small and therefore difficult to see.

**Figure 5:**
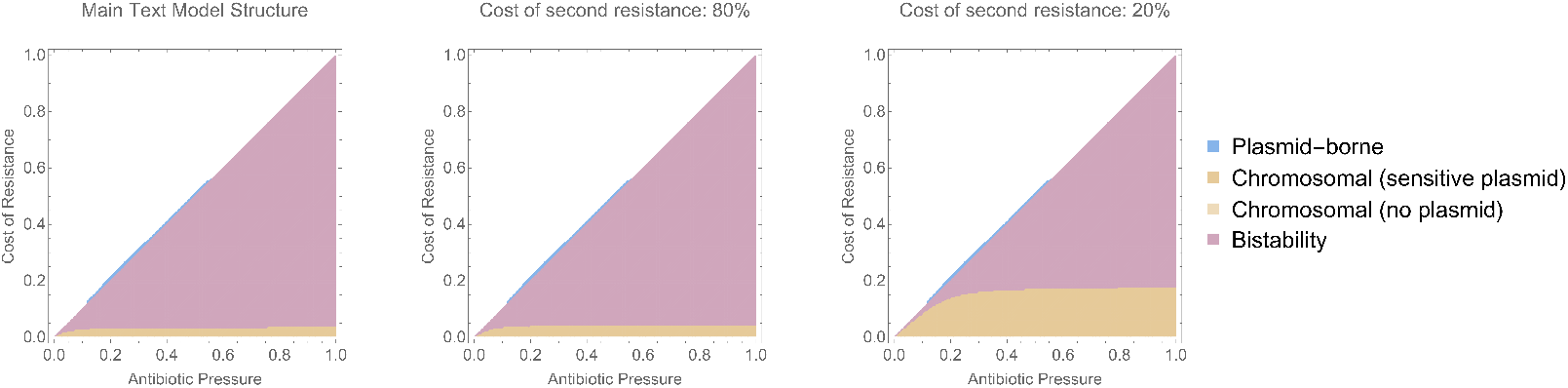
Effect of decreasing the cost incurred from a second copy of the resistance gene. Each panel shows parameter regions in which plasmid-borne only (blue); chromosomal resistance (orange); or both (purple) are evolutionarily stable, as a function of antibiotic consumption and the cost of resistance. Left-hand panel: *e* = 1 (main text model); middle panel: *e* = 0.8; right-hand panel: *e* = 0.2. Other parameters are: *λ* = 1, *γ* = 1, *s* = 0.005, *c*_*P*_ = 0.075, and *β* = 0.1. Note that the value of *β* is different from the value in the main text (*β* = 0.2) to better illustrate the region of chromosomal resistance only at low cost of resistance. With *e* = 0, resistance occurs on both chromosome and plasmid.

**Figure 6:**
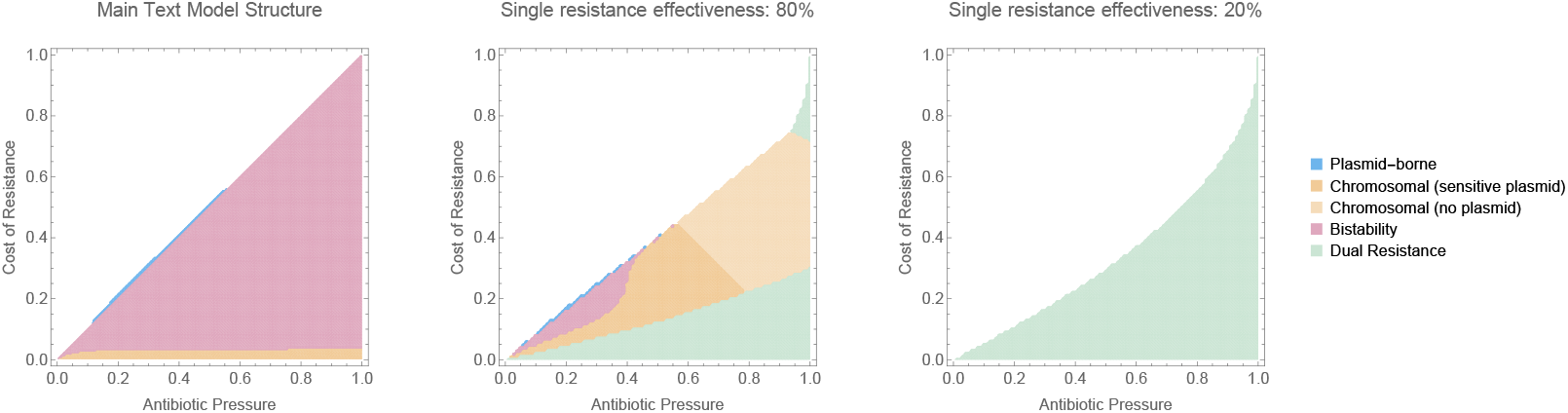
Effect of assuming a single copy of the resistance does not fully prevent antibiotic-associated mortality. Each panel shows parameter regions in which plasmid-borne only (blue); chromosomal resistance (orange); or both (purple) are evolutionarily stable, as a function of antibiotic consumption and the cost of resistance. Left-hand panel: *e* = 1 (main text model); middle panel: *e* = 0.8; right-hand panel: *e* = 0.2. Other parameters are: *λ* = 1, *γ* = 1, *s* = 0.005, *c*_*P*_ = 0.075, and *β* = 0.1. Note that the value of *β* is different from the value in the main text (*β* = 0.2) to better illustrate the region of chromosomal resistance only at low cost of resistance.

**Figure 7:**
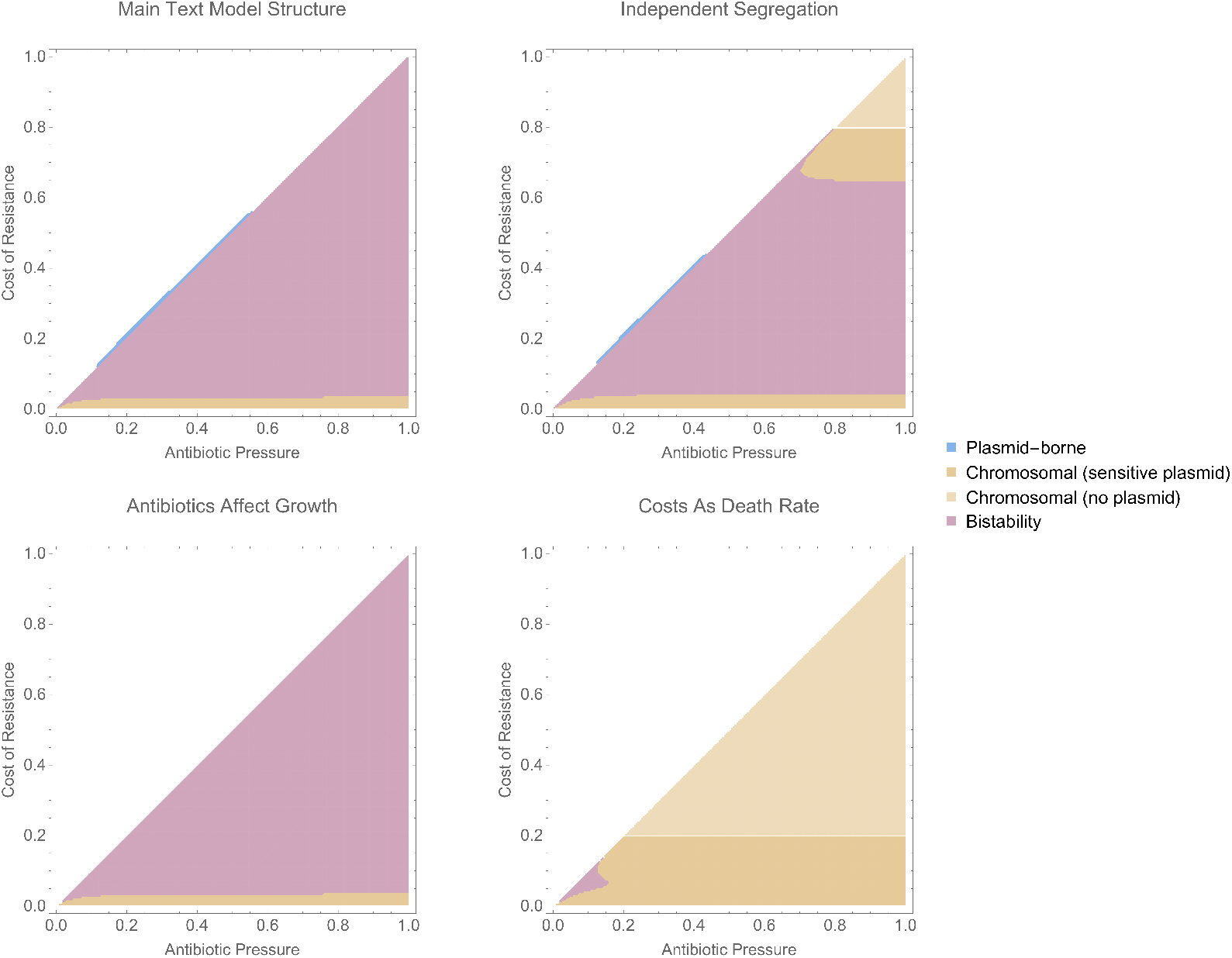
Robustness of model results to assumptions about segregation loss, the effect of antibiotic action and how fitness costs are modelled. Although the predicted outcome at specific parameter values depends on model structure, the key result that there is an area of bi-stability where either plasmid-borne or chromosomal resistance can occur is robust. Parameters are: *λ* = 1, *γ* = 1, *s* = 0.005, *c*_*P*_ = 0.075, and *β* = 0.1. Note that the value of *β* is different from the value in the main text (*β* = 0.2) to better illustrate the region of chromosomal resistance only at low cost of resistance.

#### 2.1 Cost of second resistance gene

SI Figure 5 shows the effect of decreasing the cost incurred from the second copy of the resistance gene. The model structure remains as described by Equations 1 in the main text. However, the fitness cost of the second resistance gene is modulated by a factor *e*. Thus, the replication rate of dually resistant cells is given by: *λ*_*RRP*_ = (1—*e*c*_*R*_)(1 — *c*_*R*_)(1—*c*_*P*_)*λ*. Setting *e* = 1 recovers the model from the main text, whereas *e* = 0 means a second copy of the resistance incurs no additional fitness cost. Note that if *e* = 0, resistance will occur on both the chromosome and plasmid.

#### 2.2 Benefit of second resistance gene

SI Figure 6 shows the effect of assuming a single copy of the resistance does not fully prevent antibiotic-associated mortality. Cells with a single copy of the resistance gene (i.e. either chromosomal or plasmid-borne resistance only) are subject to an antibiotic-associated mortality of (1−*a*)*A*, where *a* parametrises the effectiveness of a single resistance gene. Setting *a* = 1 recovers the model from the main text, whereas *a* = 0 would mean a single copy of the gene does not decrease antibiotic associated mortality at all. The dynamics of this modified model (with changes from the main text highlighted in bold) are described by:

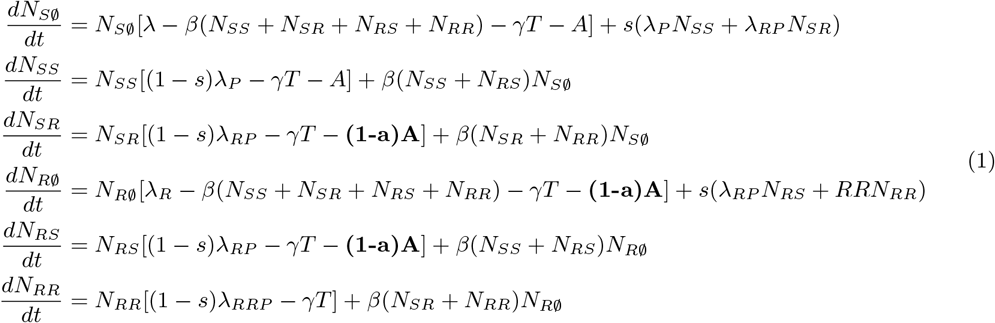

#### 2.3 Other features of model structure

SI Figure 7 explores the robustness of our results to other aspects of model structure. Although the predicted outcome at specific parameter values depends on model structure, the key result, that there is a parameter region of bi-stability where either plasmid-borne or chromosomal resistance can occur, is robust.

##### 2.3.1 Independent segregation loss

In the main text, segregation loss is modelled as occurring during cell replication. Some previous models addressing similar topics (Svara & Rankin [main text reference 6] and Bergstrom et al. [main text ref 16]) have modelled segregation loss as occurring independently of cell replication. We therefore verify that our results are robust to modelling segregation loss in this manner. The dynamics of the modified model (with the difference from the main text highlighted in bold) are given by:

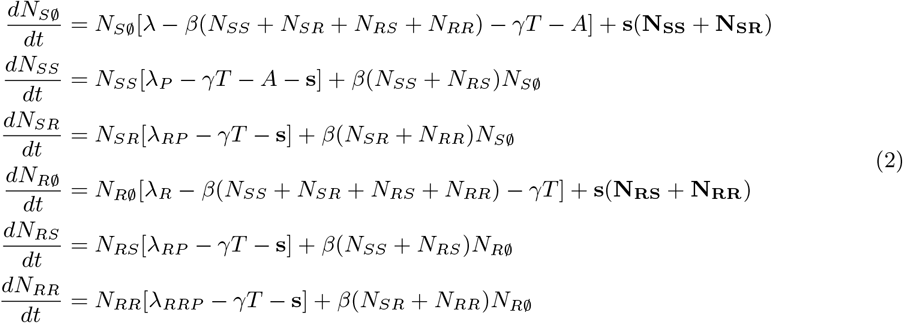

Our key result, the presence of bi-stability, is robust to this change in model structure. However, the relationship between the cost of resistance and evolutionary outcome is altered. In the main text model structure, bi-stability occurs once the cost of resistance is high enough. In this modified model, chromosomal stability is the only evolutionarily stable outcome at low *and high* cost of resistance, with bi-stability in between.

##### 2.3.2 Antibiotic effect in growth

In the main text, we model antibiotic action as an additional mortality rate (*bactericidal*). Antibiotics can also have the effect stopping cell replication (*bacteriostatic*). To test the robustness of our results to mechanism of antibiotic action, we modify the effect of antibiotics so that they decrease growth rate. The model structure remains otherwise similar to Equations 1 in the main text, but the growth rate of antibiotic sensitive cells is decreased by a factor of 1 − *A*. Thus the growth rate of *S* cells, previously *λ*, becomes (1 − *A*)*λ*, and the growth rate of *SS* cells, previously *λ*_*P*_, becomes (1 − *A*)*λ*_*P*_.

This change eliminates the region in which plasmid-borne resistance is stable but chromosomal resistance is not, but does not otherwise impact our results. Thus, our main findings apply to both bacteriostatic and bactericidal antibiotics. More generally, this broadens the relevance of our results from antibiotic resistance genes to any gene with effects on either the growth or death rate of a cell.

##### 2.3.3 Additive costs

In the main text, we model the fitness cost of resistance as a multiplicative factor on the growth rate. Here, we modify the model so that the cost of antibiotic resistance and plasmid carriage is modelled as an additional death rate. They dynamics of this modified system are described by the following equations (differences from the main text are highlighted):

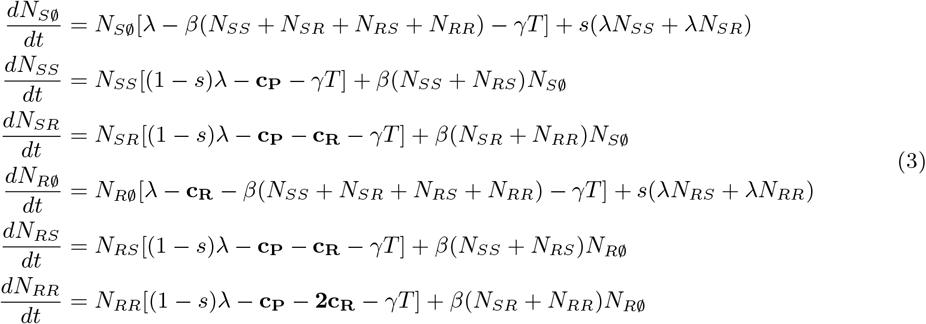

This change also eliminates the region in which plasmid-borne resistance is stable but chromosomal resistance is not. The region of bi-stability is still observed, although for this parametrisation, the region is considerably smaller. However, it should be noted that although the parameter values are the same as in the main text model, the parametrisations are no longer directly comparable because the scaling of the fitness cost is no longer the same: in the main text, fitness cost is constrained to be between 0 and 1, whereas here, the value is constrained but its effective magnitude depends on the replication and death rates (i.e. a cost of *c*_*R*_ = 0.5 for example has different impact if *λ* = 0.5 than if *λ* = 2).

### 2.4 Gene flow between plasmid and chromosome

To model the effect of genes on transposons moving between the plasmid and chromosome, we modify the model (main text Equations 1) to include a transposition term. We model replicative (‘copy-paste’) transposition: *SR* cells convert to *RR* cells at rate *ρλ*_*PR*_, and *RS* cells convert to *RR* cells at rate *ρλ*_*PR*_. The transposition rate *ρ* is scaled with replication rate to allow easy comparison of modelled transposition rates to empirical estimates expressed in terms of transposition events per generations (see below). The dynamics of the modified system are captured by:

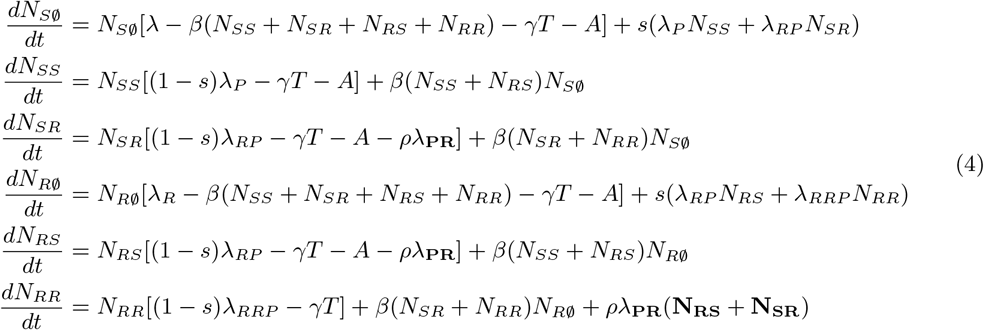

Inclusion of gene transfer between the plasmid and chromosome allows the two forms of resistance to coexist, with the frequency of the low frequency form increasing with the rate of transposition (SI Figure 8). This is analogous to mutation-selection balance. As shown in SI Figure 8, the eventual outcome remains dependent on the initial conditions, with increasing transposition rate increasing the range of initial conditions leading to chromosomal resistance. At very high rates of transposition (10^−2^ per transposon per generation), this leads to chromosomal resistance being the eventual outcome even when the initial frequency of plasmid-borne resistance is very high. However, such high rates are implausible: rate estimates for *Escherichia coli* are of the order of 10^−5^ events per generation per transposon for replicative transposition (and 10^−8^ for non-replicative transposition) [Sousa et al., main text reference 29].

**Figure 8:**
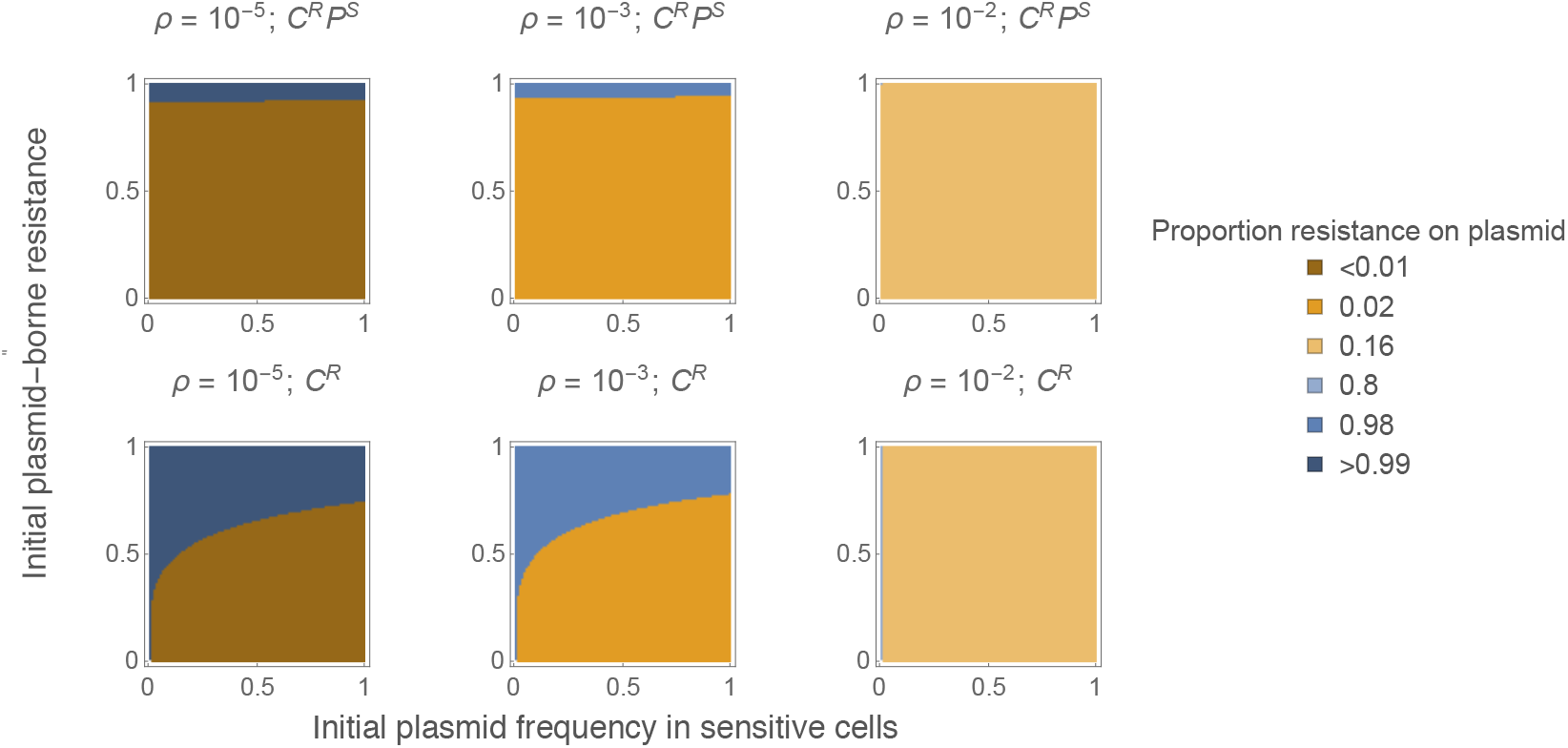
Effect of gene flow between plasmid and chromosome. The panels illustrates the proportion of resistance that is plasmid-borne at equilibrium ((*N*_*SR*_ + *N*_*RR*_)*/*(*N*_*R*∅_ + *N*_*SR*_ + 2*N*_*RR*_)). *ρ* is the transposition rate per cell per generation (for *Escherichia coli*, estimates of this rate are of the order of 10^−5^ [Sousa et al., main text reference 29]. Similarly to main text Figure 3, the x-axis indicates the frequency of the sensitive plasmid in the initial sensitive population *N*_*SS*_*/*(*N*_*S*∅_+*N*_*SS*_). The y-axis indicates the frequency of the plasmid-borne resistance in the initial resistant population 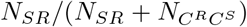 for the top panel, *N*_*SR*_*/*(*N*_*SR*_ + *N*_*R*∅_) for bottom panel). The total densities of the initial sensitive and resistant populations are both 1. Parameter values are: *λ* = 1, *γ* = 1, *s* = 0.005, *c*_*P*_ = 0.075, *β* = 0.2, *A* = 1 and *c*_*R*_ = 0.05.

#### 2.5 Imperfect exclusion of plasmid co-infection

In the main text, we assume that carrying one form of the plasmid prevents infection with the other. As such exclusion may not be fully effective, we test the effect of relaxing this assumption. For simplicity, we do not model the co-infected state explicitly: we assume that upon division of a co-infected cell, each daughter cell inherits only one of the plasmids, with equal probability of inheriting either (i.e. half of co-infections result in the super-infecting plasmid replacing the resident plasmid). With *k* indicating the probability of transferring a plasmid to a cell already carrying a plasmid (compared to a plasmid-free host cell), imperfect exclusion can be approximated as:

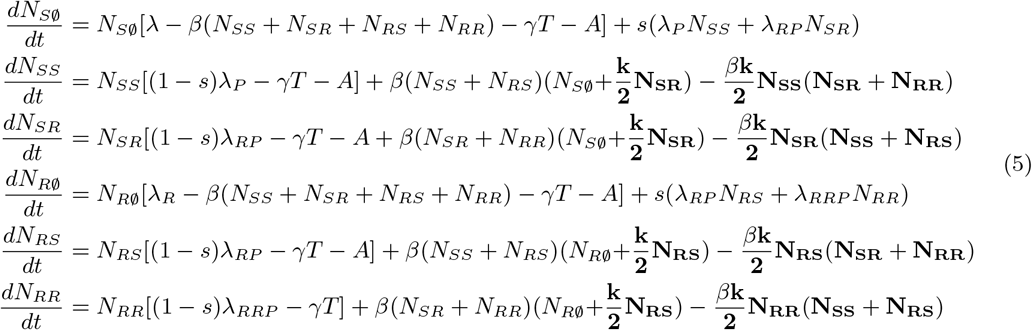

This change has no impact on the evolutionary stability of plasmid-borne and chromosomal resistance. However, within the region of bi-stability, increasing *k* increases the range of initial conditions in which the outcome is plasmid-borne resistance (SI Figure 9). This is because co-infection increases the advantage the resistant plasmid acquires from plasmid transmission, thus allowing it to increase in frequency more rapidly even in presence of the sensitive plasmid.

**Figure 9:**
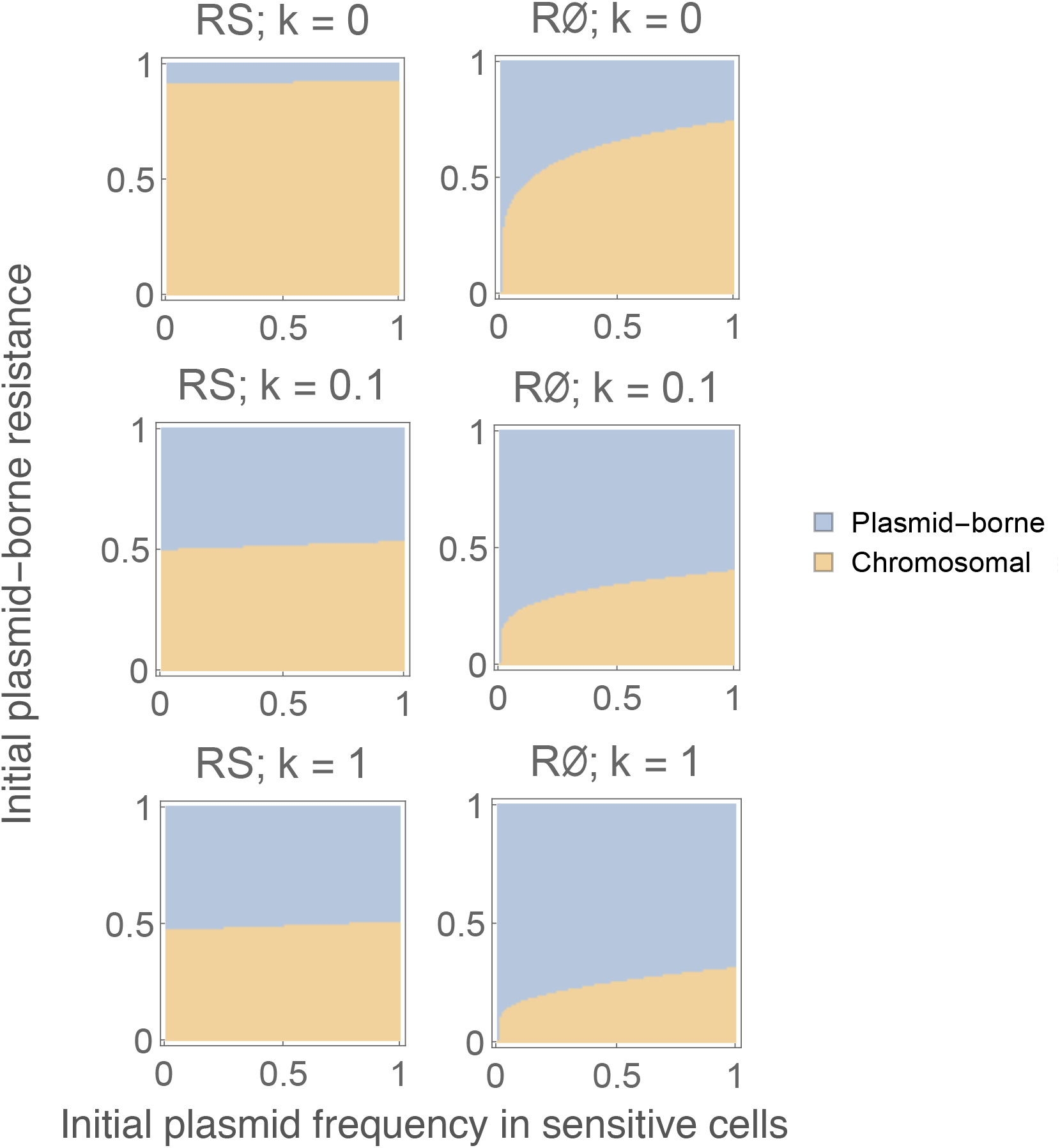
Effect of initial conditions on equilibrium location of the resistance gene, with imperfect exclusion of plasmid co-infection. As in main text Figure 3, the panels illustrates whether plasmid-borne (blue) or chromosomal resistance (orange) are observed at equilibrium. The x-axis indicates the frequency of the sensitive plasmid in the initial sensitive population. The y-axis indicates the frequency of the plasmid-borne resistance in the initial resistant population. The initial resistant population consists either of *RS* and *SR* cells (left) or *R*∅ and *SR* cells (right). The rows correspond to increasing probability of co-infection (*k*). Parameter values are: *λ* = 1, *γ* = 1, *s* = 0.005, *c*_*P*_ = 0.075, *β* = 0.2, *A* = 1 and *c*_*R*_ = 0.05.

### 2.6 Temporally fluctuating selection

In the main text, we treat the antibiotic-associated death rate as a constant. In natural settings, selection pressure from antibiotics is often heterogeneous, either through spatial (see ‘Local Adaptation’ section below) or temporal variation in antibiotic concentration. We therefore modify the model to include a fluctuating antibiotic-associated death rate. The model structure remains identical to that presented in the main text, but with antibiotic-associated death rate modelled as: *A*[1 + *sin*(2*π/T*)]. That is, a sine wave with period *T* and mean and amplitude *A*.

Because of the switch to fluctuating selection, we can no longer apply the linear stability analysis approach used in the main text to determine the evolutionary stability of plasmid-borne and chromosomal resistance in this system. We therefore use simulation to check whether established plasmid-borne resistance can be displaced by introduction of chromosomal resistance at low frequency, and vice versa.

As illustrated in SI Figure 10, the presence of bi-stability is robust to the inclusion of fluctuation. Fluctuation favours plasmid-borne resistance, but the effect is dependent on the period of the fluctuation. This relationship arises because the relative fitness of chromosomal and plasmid-borne resistance is dependent on both the relative frequencies of the two forms of resistance and the frequency of the sensitive plasmid. These frequencies are affected differently by phases of positive and negative selection. Thus, the relationship between the characteristics of fluctuating selection (i.e. phase, amplitude, period) and which resistance is favoured is likely to be complex.

**Figure 10:**
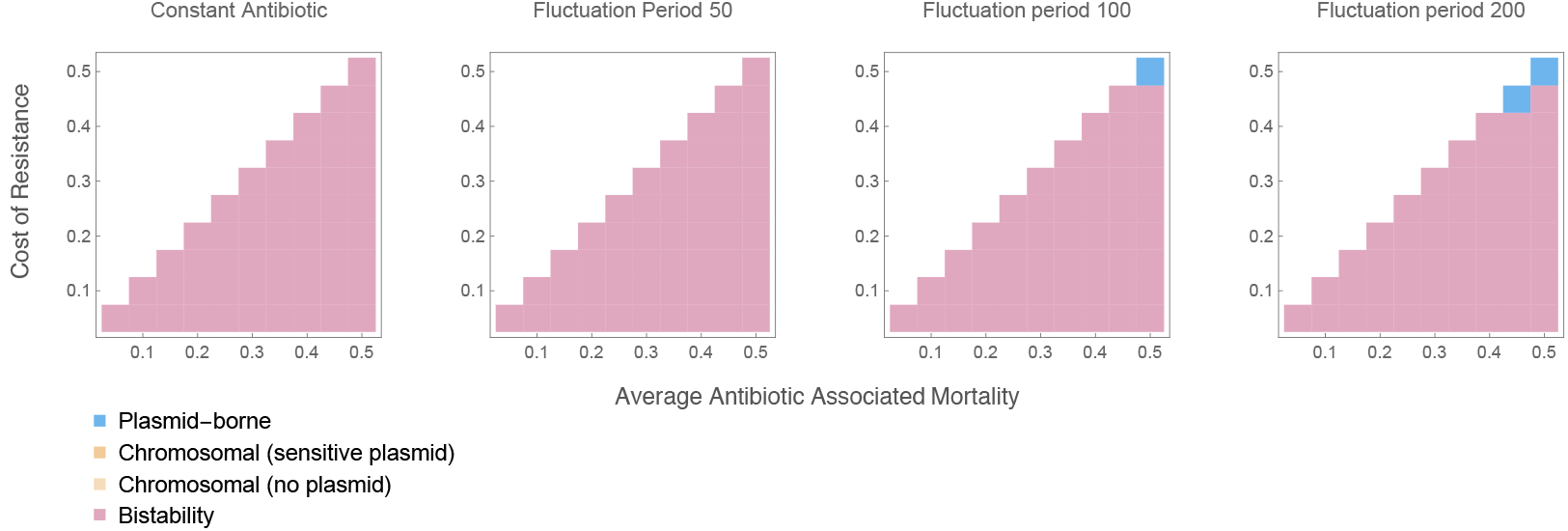
The effect of fluctuating selection on the evolutionary stability of chromosomal and plasmid-borne resistance: the presence of bi-stability is robust, with plasmid-borne resistance favoured under some conditions. Each simulation is started from the equilibrium reached when only one form of resistance (plasmid-borne or chromosomal) is included in the model. The other form is then introduced at low frequency (10^−4^). If the low frequency form of resistance is able to increase in frequency and invade, the established resistance is not evolutionarily stable. Parameter values are: *λ* = 1, *γ* = 1, *s* = 0.005, *c*_*P*_ = 0.075, *β* = 0.2, *A* = 1 and *c*_*R*_ = 0.05.

## 3 Local adaptation

We modify the model in the main text to include influx of cells from a sensitive population. Here, *μ* is the overall rate of influx of sensitive cells, and *q* is the proportion of the incoming cells which carry the sensitive plasmid. In our simulations, the initial population is fully resistant (and comprised of *SR* and *RS* cells in varying proportions). The modified dynamics are given by:

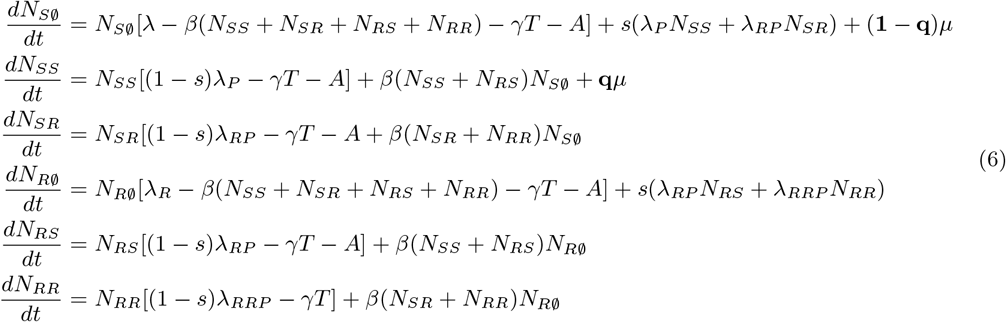

The effect of this modification depends on the frequency of the sensitive plasmid in the incoming cells (*q*): an influx of sensitive cells carrying the sensitive plasmid favours chromosomal resistance whereas an influx of sensitive cells without the plasmid favours plasmid-borne resistance (see the main text).

## 4 Plasmid persistence

We consider the relationship between plasmid transmissibility and the location of resistance genes in more detail. This relationship is relevant to the question of plasmid persistence: under the low transmissibility assumption (i.e. plasmids are not transmissible enough to persist as parasites), Bergstrom et al. show that beneficial genes will always locate on the chromosome rather than plasmid (in absence of local adaptation) [main text reference 16]. Thus, the persistence of low transmissibility plasmids is a paradox: they cannot be maintained without beneficial genes, but beneficial genes cannot be maintained on these plasmids. Here, it is worth noting that Bergstrom et al. refer specifically to persistence of a non-beneficial plasmid in a host population where the chromosome also lacks the beneficial gene. In our framing, this corresponds to persistence of the sensitive plasmid in a chromosomally sensitive population.

To check whether this result also holds in our model structure, we use linear stability analysis to compare the parameter space in which plasmid-borne resistance is evolutionarily stable with the parameter space in which parasitic plasmids are viable. In other words, we compare the space in which resistance genes can locate on the plasmid (despite competition from chromosomal resistance) with the space in which a sensitive plasmid can be maintained in a chromosomally sensitive population (in absence of competition from the resistant plasmid). Contrary to Bergstrom et al., we find that resistance genes *can* locate onto the plasmid even when plasmid transmissibility is too low for parasitic plasmids to be viable (SI Figure 11). Thus, low transmissibility plasmids can theoretically persist because of the advantage they provide their host cells.

**Figure 11:**
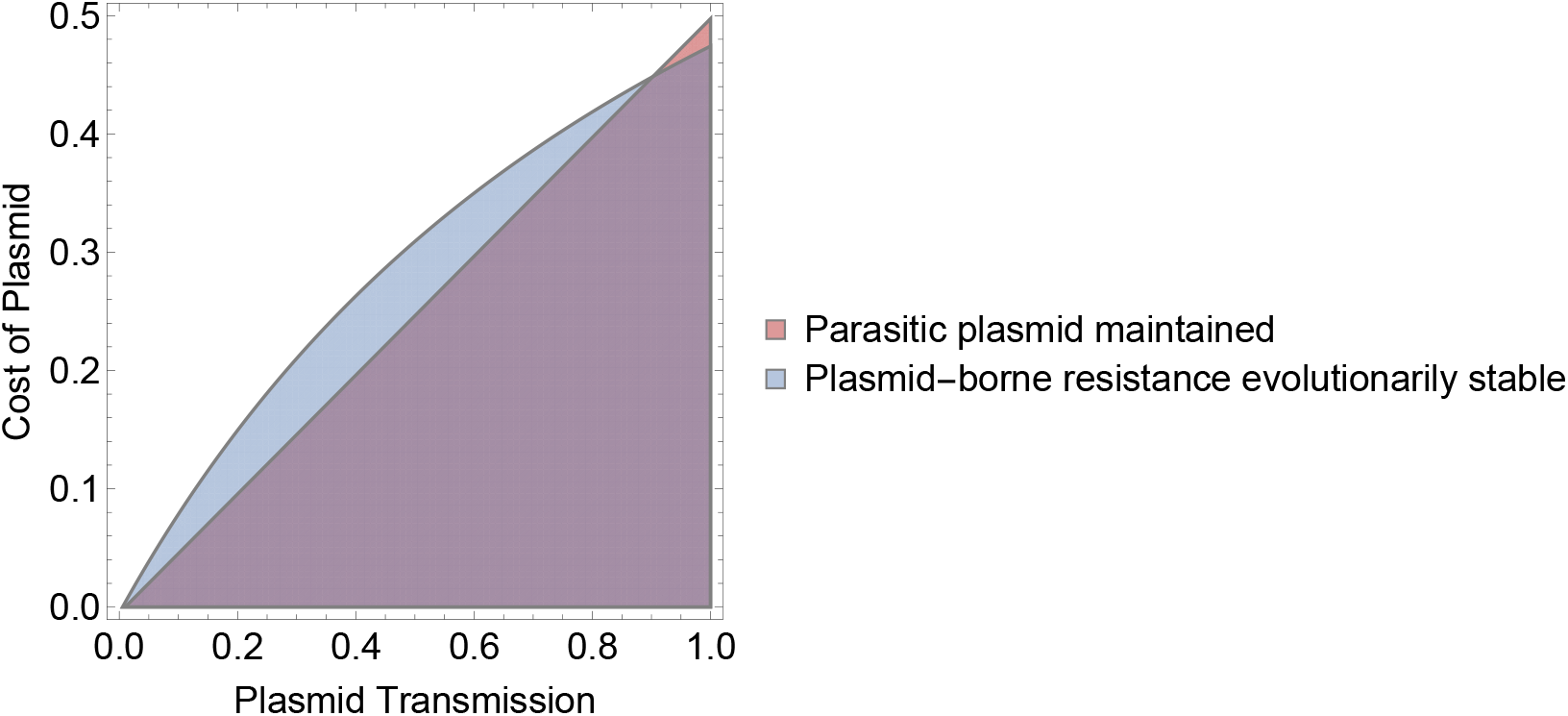
Resistance can locate onto the plasmid even when plasmid transmissibility is low, allowing low transmissibility plasmids to persist because of the advantage they provide their host cells. The plot shows the region of evolutionary stability of plasmid-borne resistance, i.e. where resistance can occur on the plasmid despite competition from chromosomal resistance (blue); the region where a parasitic plasmid (i.e. a sensitive version of the plasmid in a sensitive population) can persist (red); and the overlap of these regions (purple). Note that the blue region in which plasmid-borne resistance is evolutionarily stable is bi-stable: the resistance gene is not necessarily on the plasmid; which form of resistance is observed at equilibrium depends on the initial frequencies. Values for non-varying parameters are the same as in main text, except antibiotic associated mortality is lower (i.e. resistance is beneficial, but not essential, otherwise the sensitive cell population would not persist at all): *λ* = 1, *γ* = 1, *s* = 0.005, *c*_*P*_ = 0.075, *β* = 0.2, *A* = 0.5 and *c*_*R*_ = 0.05.

## 5 Additional Figures

**Figure 12:**
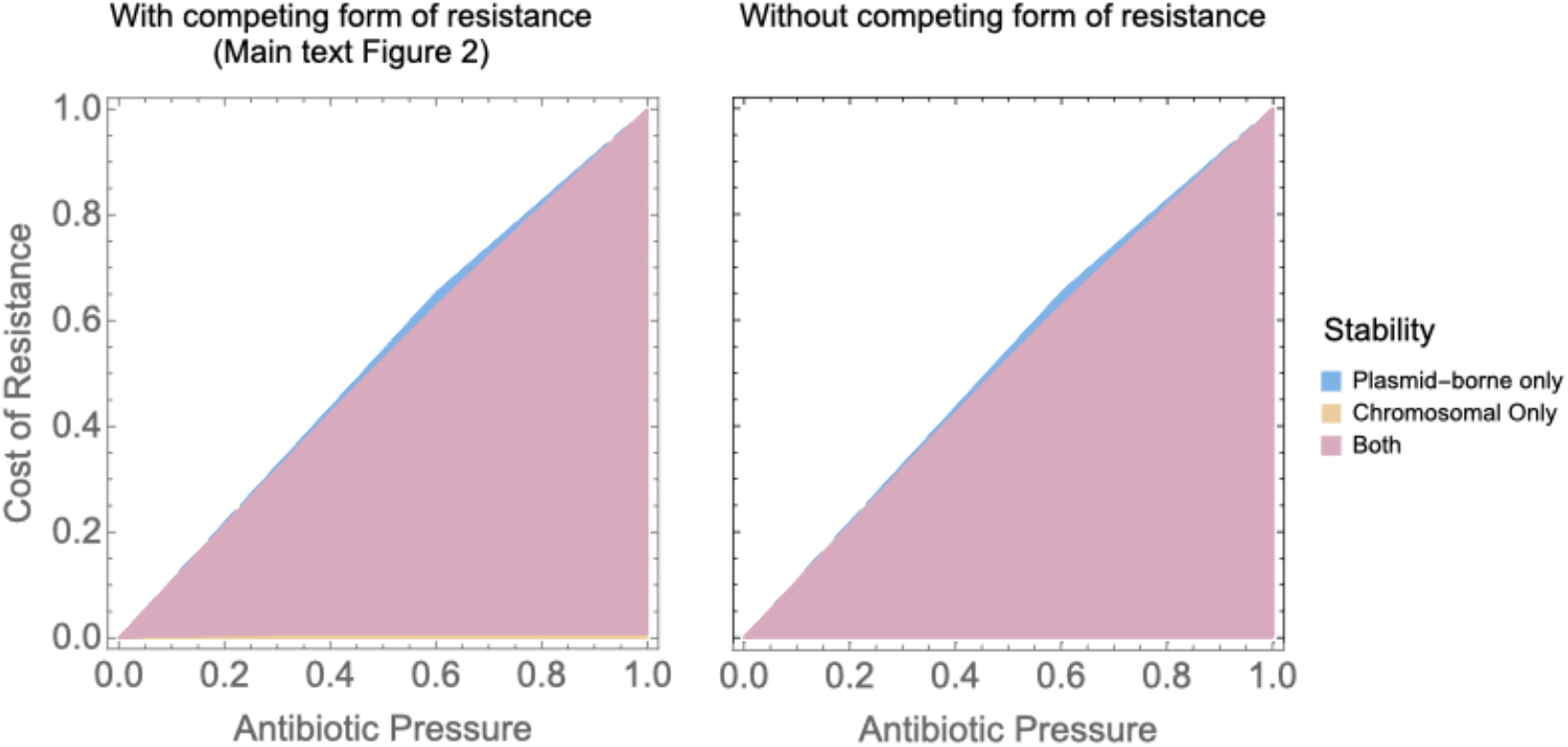
Stability of plasmid-borne and chromosomal resistance in absence of competition from the other resistance form (but in presence of the sensitive chromosome and sensitive plasmid).The left-hand panel shows the evolutionary stability depicted in main text Figure 2. The right-hand panel shows stability in absence of competition from the other resistance form. For chromosomal resistance, these two areas are the same. For plasmid-borne resistance, there is a region at low fitness cost (narrow orange band at the bottom of the left-hand panel) where plasmid-borne resistance is stable in absence of competition from chromosomal resistance, but not evolutionary stable i.e. not stable in presence of competition from chromosomal resistance. Parameter values are *λ* = 1, *γ* = 1, *s* = 0.005, *c*_*P*_ = 0.075, *β* = 0.2, *A* = 1 and *c*_*R*_ = 0.05.

**Figure 13:**
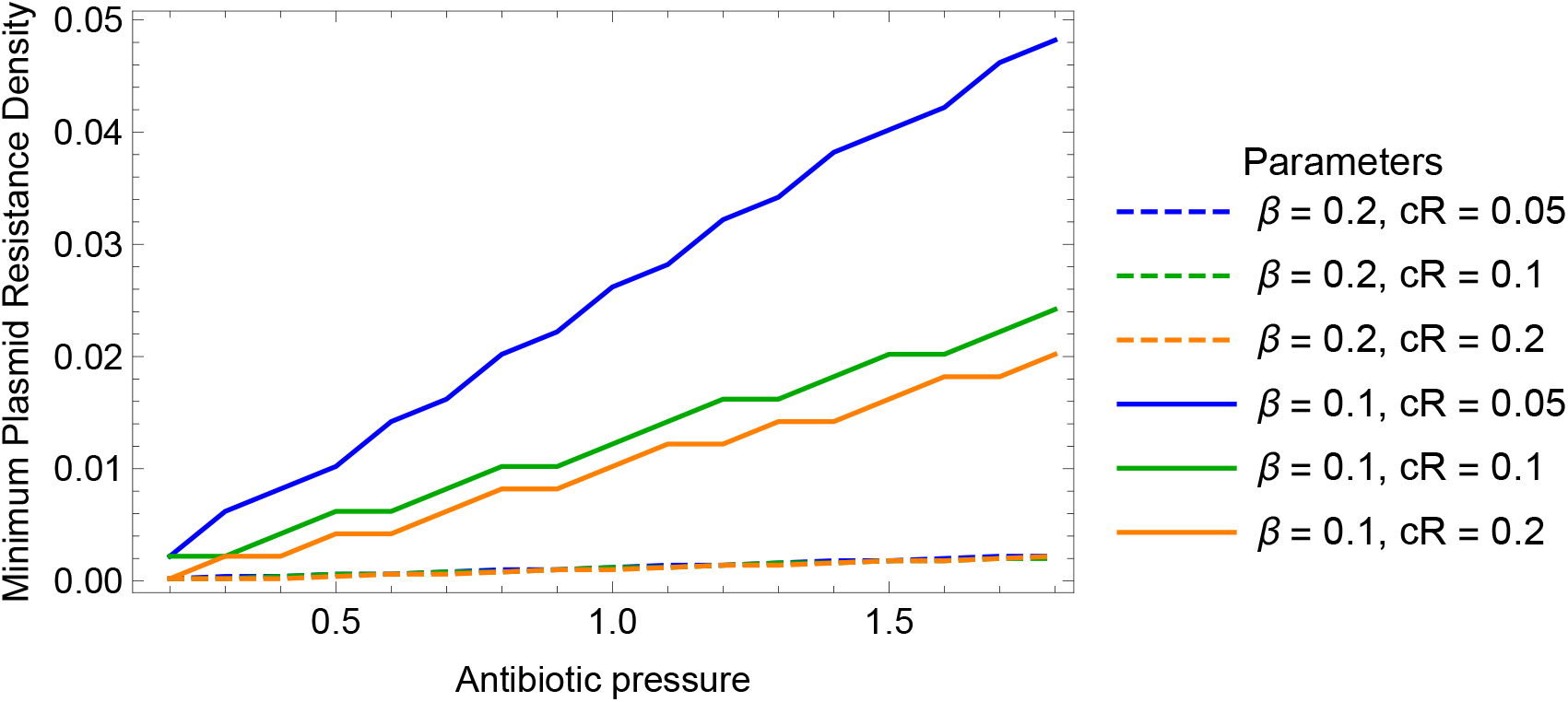
Low densities of plasmid-borne resistance are enough to prevent invasion by chromosomal resistance. The plot shows the minimum initial density of plasmid-borne resistance (*N*_*SR*_) for which chromosomal resistance introduced at low density (*N*_*RS*_ = 10^−4^) cannot invade. The initial conditions for other cell types are *N*_*S*∅_ = 0.2 and *N*_*SS*_ = 0.70 (approximating equilibrium in absence of antibiotics when *β* = 0.1), and 0 for all others. Parameters are *λ* = 1, *γ* = 1, *s* = 0.005, *c*_*P*_ = 0.075.

**Figure 14:**
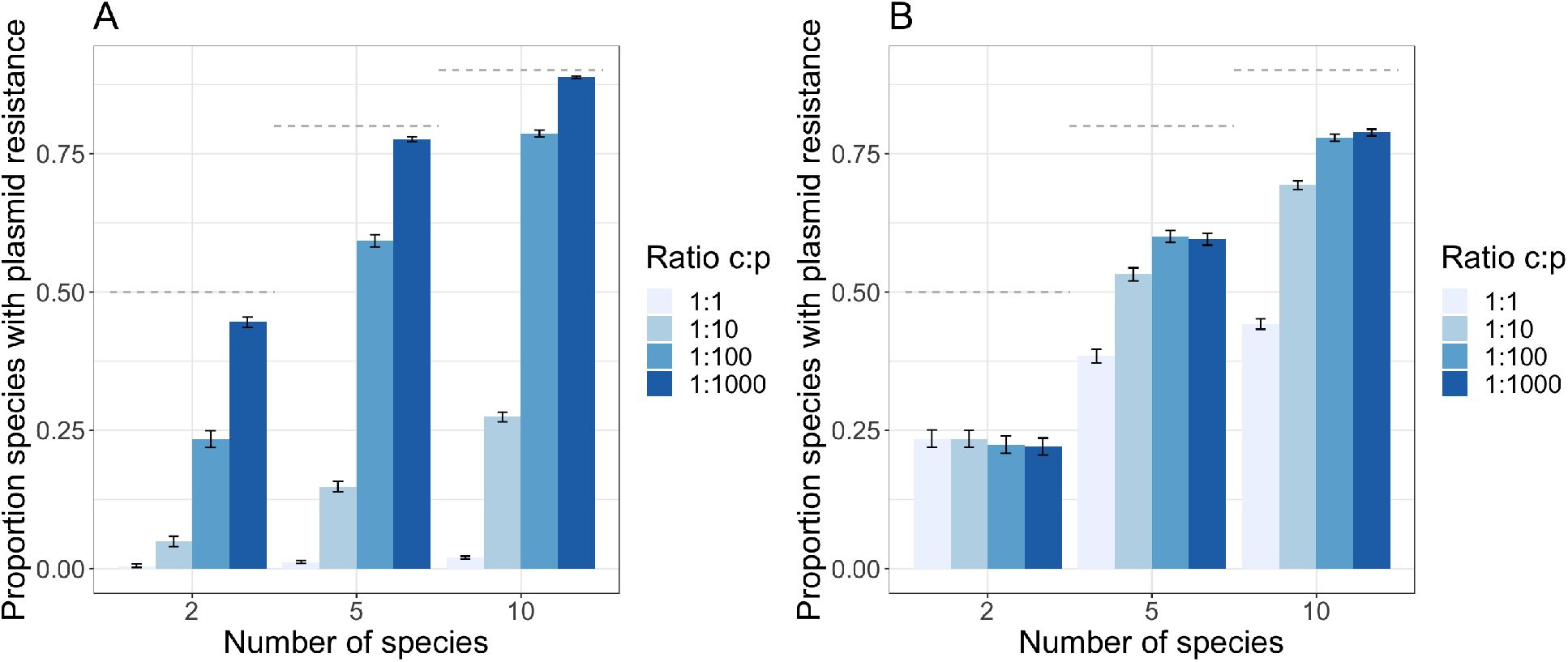
Spread of resistance genes between species for rate parameters differing from those used to generate Figure 5 in the main text. As in the main text figure, the bars indicate the proportion of plasmid-borne resistance depending on the number of simulated species and the ratio of the rate of interspecies transfer of the chromosomal (*c*) and plasmid-borne gene (*p*). The horizontal lines show the maximum proportion of plasmid resistance, given that resistance must first emerge on the chromosome ((*n*−1)*/n*). Error bars represent 95% confidence intervals based on 1000 realisations. A: With a lower rate of gene flow between gene and chromosome than in the main text (*t* = 10^−2^ instead of *t* = 10^−1^ in the main text, *m* = 10^−6^ and *c* = 10^−5^ as in the main text). The lower rate of gene flow means chromosomal resistance has more time to spread before a plasmid-borne resistance arises, leading to a lower proportion of species with plasmid borne resistance. B: Here, we assume the rate of acquiring plasmid-borne resistance from a species with chromosomal resistance is independent of the rate of interspecies plasmid transfer, and similarly, that the rate of acquiring chromosomal resistance from a species with plasmid-borne resistance is independent of the rate of interspecies transfer of chromosomal genes. In this simulation, both these rates are 10^−5^ and *m* = 10^−6^ and *c* = 10^−5^ as in the main text. The effect of variation in *p* is reduced compared to the main text because the the rate at which plasmid-borne resistance arises does not depend on *p*.

## Notes

### Competing Interest Statement

The authors have declared no competing interest.

